# A Comprehensive Structural and Functional Analysis of *Saccharomyces* Killer Toxins

**DOI:** 10.1101/2025.11.04.686668

**Authors:** Jack W. Creagh, David C. Reetz, Lily L. Givens, Sarah A. Coss, Rodolfo Bizarria, Jagdish Suresh Patel, Andre Rodrigues, F. Marty Ytreberg, Paul A. Rowley

## Abstract

Antifungal killer toxins are cytotoxic proteins that have the potential to combat the growing threat of fungi to human health and agriculture. A lack of empirical tertiary structures has placed limitations on understanding their mechanisms of action and targeting of pathogens. AlphaFold and molecular dynamics simulations were used to create tertiary structure models of all canonical *Saccharomyces* killer toxins. These models have enabled the prediction of the functional domains of killer toxins and postranslational modifications, including sites of proteolytic cleavage and disulfide bonds. They have also revealed unexpected homology between *Saccharomyces* killer toxins, suggesting that all but K28 are likely ionophores. Structural homology with the well-studied killer toxins K1 and K2 enabled prior empirical data to predict the antifungal and immunity mechanisms of the K1L, K21, K45, K74, and KHS toxins. The understudied killer toxins Klus, KHR, and K62 were found to have homology to bacterial and plant toxins, including members of the aerolysin family and antifungal lectins. These structural similarities provided clues for the mechanisms of killer toxin carbohydrate binding, oligomerization, and membrane attack. Given the hundreds of sequence homologs of the *Saccharomyces* killer toxin identified across fungi, modeling studies offer an exciting opportunity to characterize novel toxin-like proteins. This approach is strengthened by the continued use of the model yeast *S. cerevisiae* to study killer toxins and the wealth of functional data gathered in the decades since their first discovery.

## 1. Introduction

Killer yeasts produce extracellular antifungal proteins, known as killer toxins that inhibit the growth of competing yeasts and filamentous fungi [1]. Such a broad definition of a killer toxin has meant the inclusion of carbohydrate-degrading enzymes (chitinases, glucanases etc.) and nucleases, in addition to those toxins that attack cell membranes and disrupt ion homeostasis. As killer toxins are proteins, they can be considered distinct from the inhibitory properties of small molecules and peptides produced by yeasts (e.g., alcohols, mating pheromones, chelators etc.) [2–4]. The baker’s and brewer’s yeast *Saccharomyces cerevisiae* was the first species identified with antifungal activities resulting from the production of killer toxins [5]. Killer yeasts have since been identified in many fungal genera, across the Ascomycota and Basidiomycota. These yeasts are often from diverse environments, including but not limited to temperate and tropical forests, oceans, polar regions, and deserts, in association with insects, plants, fruits, and soils. Killer yeasts have also been repeatedly isolated from anthropic fermentations used for food and beverage production, as killer toxins can aid the invasion of these processes, causing spoilage. However, killer toxin production is also widely prevalent in *S. cerevisiae*, which is frequently used in commercial fermentations, particularly in wine production.

The widespread distribution of killer yeasts is thought to be driven by intense niche competition. Killer toxin production can enable killer yeasts to invade toxin-sensitive populations and increase yeast fitness under optimal conditions for toxin activity [6–12]. Exposure of yeasts to killer toxins drives the evolution of toxin resistance and is frequently observed during screens for killer yeasts, suggesting the evolution of antitoxin defense mechanisms and adaptation of cellular pathways and structures to prevent intoxication. For example, alterations in the configuration of *S. cerevisiae* cell wall mannans and beta-glucans confer killer toxin resistance by preventing toxin binding [13–18]. The killer toxin defense protein (Ktd1) also blocks K28 intoxication by mislocalization of the toxin to the vacuole [19,20]. Thus, the production of killer toxins influences microbial community composition by selecting for resistance and driving the evolution of novel toxins in an ongoing genetic arms race between yeasts and their toxins [21,22]. Indeed, signatures of positive selection have been detected in both *KTD1* and killer toxins [20,23,24]. Such evolutionary conflict could explain why killer yeasts tend to exhibit a narrow spectrum of antifungal activity limited to specific strains of closely related yeast species.

The growing threat of fungal pathogens that are devastating wild animal and plant populations, reducing agricultural yields, and impacting human health highlights the critical need for innovative antifungal strategies. The ability of killer yeasts to suppress pathogenic and spoilage fungi has generated interest in their potential use in agriculture and medicine. Specifically, killer yeasts can inhibit the growth of a wide diversity of human pathogenic fungi, such as the WHO priority species *Nakaseomyces glabratus* (syn. *Candida glabrata*) and *Candida albicans* [25–30] . Similar laboratory screens have been used to identify killer yeasts that can inhibit plant pathogens and spoilage fungi. This has led to the application of killer yeasts onto plants, fruits, and silage, to prevent disease and spoilage. Killer toxins have also been used to develop genetically modified, disease-resistant plants such as tobacco, maize, and tomato. In field trials, genetically modified maize expressing the killer toxin KP4 gained significant protection from a disease caused by smut fungi (Allen et al., 2011). Killer yeasts have also shown promise in protecting wine from spoilage yeasts and to prevent spoilage by diastatic strains of yeasts during brewing. Importantly, killer toxins are likely non-toxic to humans due to their prevalence in strains of *S. cerevisiae* used in food and beverage production and their lack of activity at physiological temperatures and pH. Additional safety studies suggest that killer toxins are also non-toxic to cultured human cells due to their targeting of cell surface receptors and cellular structures that are specific to fungi (e.g., mannans, chitin, and glucans). These findings underscore the potential of killer toxins as novel antifungals, offering promising solutions to address the rising challenges posed by fungal pathogens relevant to human health and agriculture.

Killer toxins from the *Saccharomyces* genus of yeasts are among the most well-studied, due to the powerful genetic tools that have been developed for use in the model yeast *S. cerevisiae*. The first killer toxin discovered was produced by *S. cerevisiae* and was later named K1 [5]. Attempts to characterize the gene responsible for K1 expression led to the determination that it was inherited cytoplasmically in a non-Mendelian fashion [31]. The nucleic acids responsible for the K1 killer phenotype were subsequently determined to be double-stranded RNAs (dsRNAs) [32]. In many *S. cerevisiae* killer yeast strains, “M” (Medium; ∼1,500 bp) dsRNAs are associated with toxin production and have been historically referred to as killer viruses or satellite viruses, but are now classified by the International Committee on Taxonomy of Viruses (ICTV) as dsRNA satellites. M dsRNAs are always associated with “L” (Large; ∼4,600 bp) dsRNAs [33–36]. L was identified as the dsRNA genome of viruses from the *Totiviridae* family that encodes proteins that enable the replication and packaging of both L and M dsRNAs in viral particles. Surveys of *S. cerevisiae* killer yeasts have revealed a significant correlation between the presence of L and M and the killer phenotype [37]. Advances in nucleic acid sequencing technology have enabled the identification of different types of M-encoded toxins, including K1, K2, K28, and Klus from *S. cerevisiae*, as well as K1L, K21 (also known as K66), K45, K62, and K74 from *Saccharomyces paradoxus* (Table 1). Killer toxin-encoding M satellites are also present in other yeasts of the Ascomycetes and Basidiomycetes.

**Table 1.**
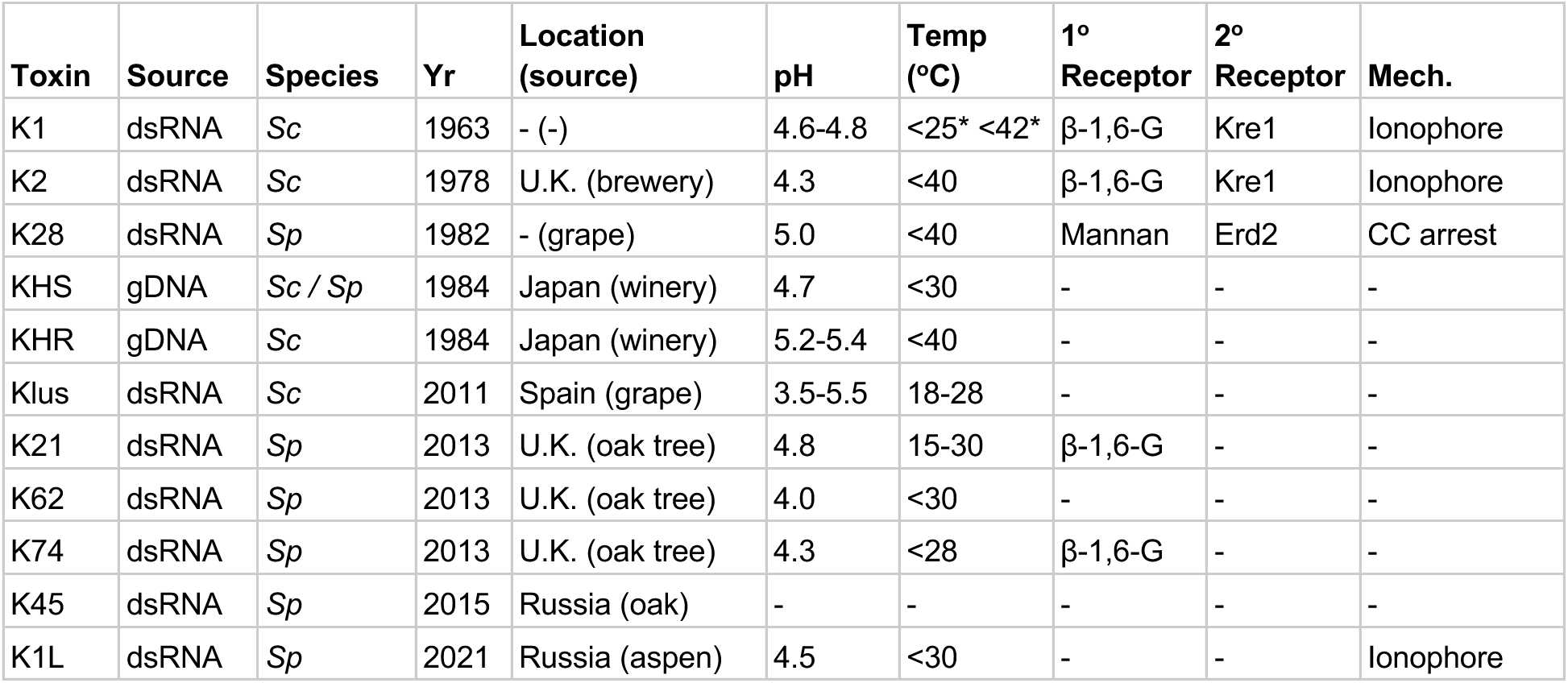
Discovery and general properties of *Saccharomyces* killer toxins. Sc, *S. cerevisiae*; Sp, *S. paradoxus*; β-1,6-G, β-1,6-glucan; ‘-’, unknown; CC arrest, cell cycle arrest; *depending on assay conditions.

| Toxin | Source | Species | Yr | Location (source) | pH | Temp (°C) | 1° Receptor | 2° Receptor | Mech. |
| --- | --- | --- | --- | --- | --- | --- | --- | --- | --- |
| K1 | dsRNA | Sc | 1963 | - (-) | 4.6-4.8 | <25* <42* | β-1,6-G | Kre1 | Ionophore |
| K2 | dsRNA | Sc | 1978 | U.K. (brewery) | 4.3 | <40 | β-1,6-G | Kre1 | Ionophore |
| K28 | dsRNA | Sp | 1982 | - (grape) | 5.0 | <40 | Mannan | Erd2 | CC arrest |
| KHS | gDNA | Sc / Sp | 1984 | Japan (winery) | 4.7 | <30 | - | - | - |
| KHR | gDNA | Sc | 1984 | Japan (winery) | 5.2-5.4 | <40 | - | - | - |
| Klus | dsRNA | Sc | 2011 | Spain (grape) | 3.5-5.5 | 18-28 | - | - | - |
| K21 | dsRNA | Sp | 2013 | U.K. (oak tree) | 4.8 | 15-30 | β-1,6-G | - | - |
| K62 | dsRNA | Sp | 2013 | U.K. (oak tree) | 4.0 | <30 | - | - | - |
| K74 | dsRNA | Sp | 2013 | U.K. (oak tree) | 4.3 | <28 | β-1,6-G | - | - |
| K45 | dsRNA | Sp | 2015 | Russia (oak) | - | - | - | - | - |
| K1L | dsRNA | Sp | 2021 | Russia (aspen) | 4.5 | <30 | - | - | Ionophore |

Despite the prevalence of toxin-encoding M dsRNA satellites in *Saccharomyces* yeasts, they appear relatively rare in killer yeasts outside of the Saccharomycotina. While there are several examples of killer toxins encoded by double-stranded linear DNA plasmids, many killer toxins are likely genome-encoded. For example, the killer toxins KHR and KHS are found in the genomes of most *S. cerevisiae* strains, as well as other *Saccharomyces* and non-*Saccharomyces* species [37–41]. Genome-encoded homologs of the M satellite-encoded killer toxins Klus and K62 have also been identified in *S. cerevisiae* [42]. Whole-genome sequencing techniques have enabled the discovery of hundreds of putative killer toxins based on primary sequence homology to known killer toxins. One prominent example is the killer toxin KP4 produced by the Basidiomycete yeast *Ustilago maydis,* which has hundreds of genome-encoded homologs in fungi, as well as non-vascular plants [43–46]. The large numbers of putative killer toxins in the KP4 family and other groups of genome-encoded killer toxins have provided evidence for the horizontal gene transfer, rapid evolution, and gene expansion of killer toxins. The widespread distribution and diversification of killer toxin homologs underscore their importance in fungal ecology and evolution, suggesting they play multifaceted roles in competition, defense, and niche adaptation across diverse taxa. However, given the large and expanding number of these genes, it is a significant challenge to elucidate if they are toxins or if they have other functional roles.

The lack of knowledge regarding the antifungal mechanisms of killer toxins is partly due to a paucity of tertiary structure models. To date, only six empirical structural models of fungal killer toxins have been determined, including those from *Millerozyma farinosa* (Salt Mediated Killer Toxin, SMKT), *Ustilago maydis* (Killer Protein 4 (KP4), and Killer Protein 6 (KP6)), *Zymoseptoria tritici* (Zt-KP6-1 and Zt-KP4-1), and *Williopsis mrakii* Killer Toxin (WmKT) (Table S1). These killer toxins are all small proteins (<223 amino acids) and, except for WmKT, have elements of structural similarity based on a typical alpha/beta sandwich. Solving the tertiary structures of *Saccharomyces* killer toxins has proved challenging due to the sometimes complex workflows required for the purification of native toxins, low levels of expression, and toxicity to yeast when overexpressed [47–53]. Powerful recombinant expression systems that have been developed for bacteria have not been utilized due to the requirement for eukaryotic-specific posttranslational modification of killer toxins (i.e., protease cleavage, disulfide bond formation, glycosylation). Recombinant systems for killer toxin expression by yeasts have been developed, but none have been successfully applied to solving the tertiary structures [54–56].

Recent advances in artificial intelligence for predicting tertiary structure have enabled the creation of millions of models from primary protein sequences. The most well-known example, AlphaFold, has revolutionized tertiary structure prediction by using deep neural networks to generate high-confidence structural models without requiring an empirical template. Afterward, molecular dynamics (MD) simulations are often employed to test the structural stability and model biophysical properties and atomic interactions. We note that by contrast, classical tertiary structure prediction tools such as MODELLER, PHYRE, and PSIPRED rely on threading models based on existing empirically determined structures with shared sequence homology. While these approaches have been used to create structural models of the KP4 family of killer toxins, they are limited by the availability of experimentally determined structures, a significant weakness for killer toxin structural prediction given the limited number of empirical models [45]. Therefore, cutting-edge computational approaches using both artificial intelligence and MD provide a powerful approach for inferring killer toxin structure, posttranslational modification, and mechanism that has been previously unattainable.

This manuscript provides a comprehensive review of all current knowledge of *Saccharomyces* killer toxin function and places it in context with tertiary structural models generated by AlphaFold and MD. The long history of empirical research into the function of killer toxins (particularly K1, K2, and K28) is used to strengthen and validate these structural models. This approach will also help to predict the structure, function, and posttranslational modifications of killer toxins that have not been extensively studied by empirical methods. Results suggest that the majority of killer toxins are ionophoric, with structural homology to other fungal killer toxins and protein toxins from bacteria and plants with known mechanisms or empirical structures. Based on primary and tertiary structure analysis, we organize *Saccharomyces* killer toxins into families that we predict have similar mechanisms of action. These models will serve as a useful framework for future empirical investigations of *Saccharomyces* killer toxins and allow the categorization of thousands of putative protein toxins found in fungi and other organisms.

## 2. Results and Discussion

To analyze the sequence similarity between *Saccharomyces* killer toxins and identify killer toxin homologs, the amino acid sequences of K1, K1L, K2, K21, K28, K45, K62, K74, KHR, KHS, and Klus were used as queries for position-specific iterated BLAST (PSI-BLAST) (Figure S1 and File S1). A total of 4,437 potential homologs were identified. Sequences were filtered to exclude those shorter than 75% or longer than 150% of the query sequence, removing 1,703 candidates likely representing non-functional genes or proteins with possible functional divergence from killer toxins (Table S2). After filtering, sequence homologs were identified for all canonical killer toxins, except for K28, which only returned three near identical sequences from different strains of *S. cerevisiae* (Figure 1A and B). Across all of the identified homologs, 96% were identified in fungi, of which 93.42% were from the Ascomycota and 2.60% from the Basidiomycota and Chytridiomycota. The majority (35.4%) of the identified homologs were in the subphylum Saccharomycotina (Figure 1A). The remainder of killer toxin homologs were identified in bacteria (1.8%) and plants (0.2%). Of the 48 bacterial homologs, 37 were homologs of K62, with the remainder similar to K45 (n = 9), K74 (n = 1), and KHR (n = 1). Killer toxin homologs in plants were confined to K62 (n = 2) and K45 (n = 1). Finding killer toxin homologs encoded in the genomes of a wide diversity of organisms supports their horizontal transfer between species, as well as ongoing gene diversification and expansion.

**Figure 1.**
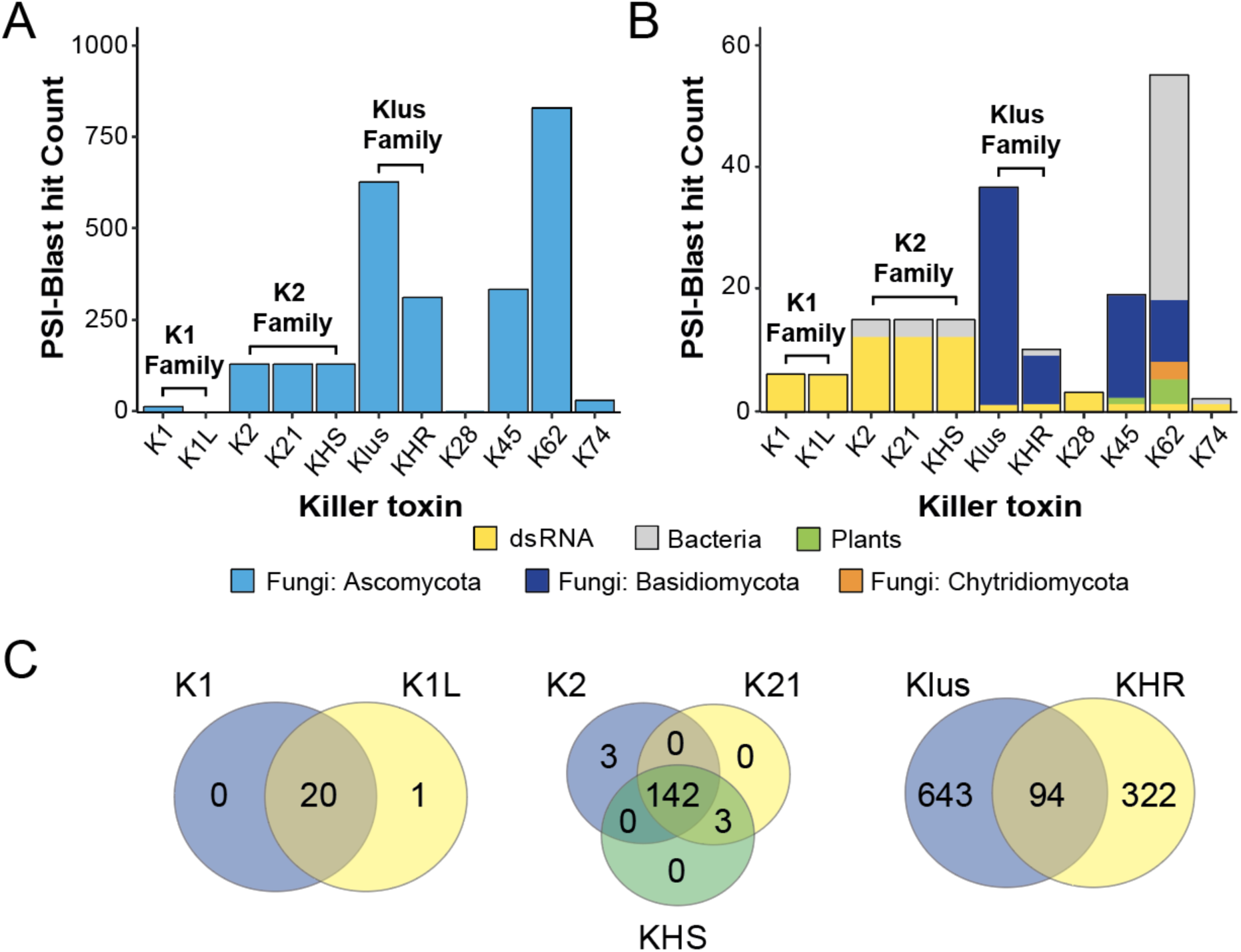
Sequence homologs of *Saccharomyces* killer toxins are abundant in fungi but are also found in plants and bacteria. (A) Identification of sequence homologs of 11 *Saccharomyces* killer toxins in the Ascomycota using the amino acid sequences of each canonical toxin to query the NCBI database using PSI-BLAST. Killer toxin families were identified based on overlap in sequence homologs between killer toxins K1/K1L (K1 family), K2/K21/KHS (K2 family), and Klus/KHR (Klus family). (B) Homologs of *Saccharomyces* killer toxins identified in fungi of the Basidiomycota and Chytridiomycota, bacteria, and plants. Killer toxins identified as being encoded on dsRNAs are also indicated. (C) Venn diagrams illustrating the overlap between sequence homologs of K1/K1L, K2/K21/KHS, and Klus/KHR.

PSI-BLAST of K1 and K1L confirmed their sequence homology and identified the majority of homologs as being from Saccharomycotina yeasts that were previously identified [23]. The 21 homologs identified by K1 and K1L showed almost perfect overlap and were collectively named the K1 family (Figure 1C). The overlap in PSI-BLAST results observed across other *Saccharomyces* killer toxins enabled the identification of two additional toxin families encompassing K2/K21/KHS (K2 family) and Klus/KHR (Klus family) (Figure 1). The sequence homology between K2 and KHS has been previously reported in other yeasts of the Saccharomycotina (e.g. *Vanderwaltozyma polyspora*) [41]. K2, K21, and KHS identified a total of 150 unique proteins, with 97.9% of the homologs shared across all three killer toxins (Figure 1C). The Klus family has the second-largest number of homologs (after K62), with 871 proteins and a relatively small overlap of just 94 proteins between Klus and KHR. PSI-BLAST analysis of the three other *Saccharomyces* killer toxins (K62, K45, and K74) identified lists of unique homologs and were named for the canonical killer toxin (Figure 1B and C).

### 2.1 Molecular Modeling of *Saccharomyces* Killer Toxins

To gain functional insights and allow the determination of structural similarities between the 11 known *Saccharomyces* killer toxins, tertiary structures were predicted with AlphaFold2 [57]. AlphaFold2 estimated the confidence of each tertiary structure by predicted Local Distance Difference Test (pLDDT) for each residue of each killer toxin. Scores can range from the lowest (0) to the highest confidence (100). Values greater than 80.0 are considered of high confidence and most often correlate with regions of secondary structure, while lower scores indicate less certain structural predictions and regions of protein disorder.

The average pLDDT for all killer toxin models ranged from 35.0 to 82.0, with K62 being the most confident and K28 being the least (Table S3, Figure S2). The pLDDT score of K74 was also low (35.1). The majority of the killer toxin models (K1, K1L, K2, K21, Klus, KHR, and KHS) had moderate confidence, with average pLDDT scores ranging from 51.0 to 68.0. Despite lower global scores, all of these models had elements of secondary structure with high confidence and an average pLDDT maximum of 89.1. AlphaFold3 was also used to generate models of each killer toxin. However, only three of the 11 AlphaFold3 models of killer toxins had improved pLDDT average scores compared to AlphaFold2. These models were KHS, K62, and Klus, which improved by pLDDT 7.5, 0.9, and 11.6, respectively. Since AlphaFold2 resulted in higher average pLDDT scores these models were used for further analyses.

AlphaFold2 structures were subjected to a 1 μs MD simulation to model their behavior in a solvated environment and to improve the quality of the predicted structures. For each toxin, the resulting trajectory from MD was clustered using the GROMACS clustering tool based on pairwise Root Mean Squared Deviation (RMSD). The central structure of the largest cluster was chosen as the final tertiary structure model for each killer toxin. After MD simulation using the initial AlphaFold2 tertiary model, the RMSD values were used to assess structural stability over time by comparing the movement of polypeptide backbone alpha carbons. Low RMSD values meant little change from the initial AlphaFold2 tertiary mode during MD simulation. In contrast, high RMSD values indicated movement of the tertiary and secondary structure away from the initial model. RMSD values that did not change significantly during MD indicated that a structure had adopted a stable conformation, even if there were initial changes in the RMSD. Structures with RMSD values that varied during MD indicated protein models that were more flexible, dynamic, or unstable.

Ramachandran plots before and after MD were used to visualize phi and psi bond angle improvements, and provided a graphical representation of favored and unfavored angles of all amino acid residues in each tertiary structure model. Folded proteins typically have most of their residues in favored regions of the plot, corresponding to elements of secondary structures, such as alpha-helices and beta-sheets. Overall, MD simulations refined the predicted killer toxin models by allowing backbone flexibility, resulting in an increased number of residues in favored regions and enhanced model quality.

### 2.2 Measurement of tertiary structure model confidence

RMSD over time for most killer toxin tertiary models stabilized between 0.4 and 2.4 nm from the initial AlphaFold structure, indicating only small structural fluctuations (Figure S2). Most models stabilized within 100 to 200 ns, with K1, K2, KHS, and K45 stabilizing quickly and showing minimal RMSD movement for the remainder of the simulation. In contrast, KHR, Klus, and K62 each had a shift in RMSD during simulation, likely due to a conformational change. For K62, this shift occurred within 50 ns, stabilized after 100 ns, and was primarily due to N-terminal flexibility. For Klus and KHR, the shift occurred after 600 ns due to flexibility in the first 15 N-terminal residues and a large flexible loop (amino acids 111-161), respectively. For K62, Klus, and KHR, removal of these flexible regions resulted in more stable structures (Figure S2). The RMSD of K1L and K28 both increased throughout the simulation, indicating the general instability of these models, likely due to the lower-confidence AlphaFold models used before the MD simulation. Overall, MD improved Ramachandran-favored residues by 3.2% on average and reduced outliers by 2.3% (Figure S2 and Table S3). These optimized models aim to place decades of empirical data on the functionality of *Saccharomyces* killer toxins in the context of protein tertiary structure.

### 2.3 The K1 killer toxin family

K1 was the first killer toxin discovered and is one of the best understood due to decades of empirical investigation. K1 was also the first killer toxin to be identified as being encoded on a dsRNA satellite (M1) associated with a totivirus. The location of the first isolation of a K1 killer yeast is unknown, but K1 is produced by many strains of *S. cerevisiae* isolated worldwide. Variants of K1 containing non-synonymous polymorphisms have also been reported to exhibit different antifungal activities [37,58]. Overall, K1 is a potent antifungal toxin that inhibits many strains and species of yeasts and is particularly effective against the opportunistic human pathogen *N. glabratus* [26,28] . The toxin is heat-labile, with an optimal temperature for activity being ∼25°C. Like many killer toxins, K1 is most active in acidic conditions (Table 1).

K1L was identified as being produced by *S. paradoxus* Y-63717, which was originally isolated from the exudate of an Asian aspen tree (*Populus davidiana*) in the eastern province of Khasan, Russia (ARS Culture Collection (NRRL)). K1L is encoded on a dsRNA satellite named M1L that is maintained by a totivirus (L-A-45) that was also found to support the replication of M45 (encoding K45) in a different strain of *S. paradoxus* from Eastern Asia [23]. The K1L toxin was identified as similar to K1 by predicted secondary structure and apparent domain organization, despite only 18% identity to K1. K1L is more closely related to a group of genome-encoded homologs found across yeasts of the Saccharomycotina, which are active killer toxins when ectopically expressed by *S. cerevisiae* [23]. K1L has a unique spectrum of antifungal activity that is more similar to K1 than other *Saccharomyces* killer toxins, which allowed its initial identification as a novel killer toxin. However, K1L is less effective at inhibiting yeast growth than K1 but is also heat-labile and has a pH optimum of 4.5 [23].

#### 2.3.1 The K1 killer toxin family: Domain organization and posttranslational modification

K1 is divided into four domains: delta (amino acids 1-44), alpha (45-147), gamma (148-233), and beta (234-316) from the N-terminus (Figure 2A). These domains are defined by sites of proteolytic processing during maturation of the 35 kDa (316 amino acids) primary translation product of K1, known as the preprocessed toxin (ppTox) [59]. To enter the secretory pathway, ppTox is exported to the endoplasmic reticulum (ER), where an N-terminal signal sequence in the delta domain is cleaved after residue A26, forming the protoxin (pTox) [60,61]. The pTox is glycosylated, although mutation of residues targeted for glycosylation has no appreciable effect on the killer phenotype. The K1 sequence contains six cysteine residues with a predicted interdomain disulfide bond between C92 and C239 of the alpha and beta domains, respectively [62]. Additional intradomain disulfide bonds are predicted between C95-C107 and C248-C312 based on their importance for immunity and toxicity [62]. The glycosylation and crosslinking of pTox results in a ∼43 kDa pTox. The enzymatic removal of carbohydrates or chemical blocking of glycosylation reverts pTox to ∼33 kDa, which is the expected molecular weight of ppTox minus the signal sequence [63,64]. Export of pTox to the Golgi is dependent on proteins of the secretory pathway, as blocking Golgi trafficking prevents further maturation of K1 by proteolytic processing [64]. Cleavage of K1 pTox occurs by the action of the Kex2 endopeptidase after basic residues R44, R149, and R233 [65,66]. A third potential Kex2 cleavage site is located after R188 in the gamma domain, and mutation of this site reduces K1 toxicity but not immunity [67]. After Kex2 cleavage, Kex1 carboxypeptidase cleaves before R148 to remove an arginine dipeptide, creating the final C-terminal end of the alpha domain [65,68]. Therefore, the mature 20.6 kDa K1 is a disulfide-linked heterodimer of processed alpha (11.1 kDa) and beta (9.5 kDa) domains linked by a single disulfide bond between C92 and C239.

**Figure 2.**
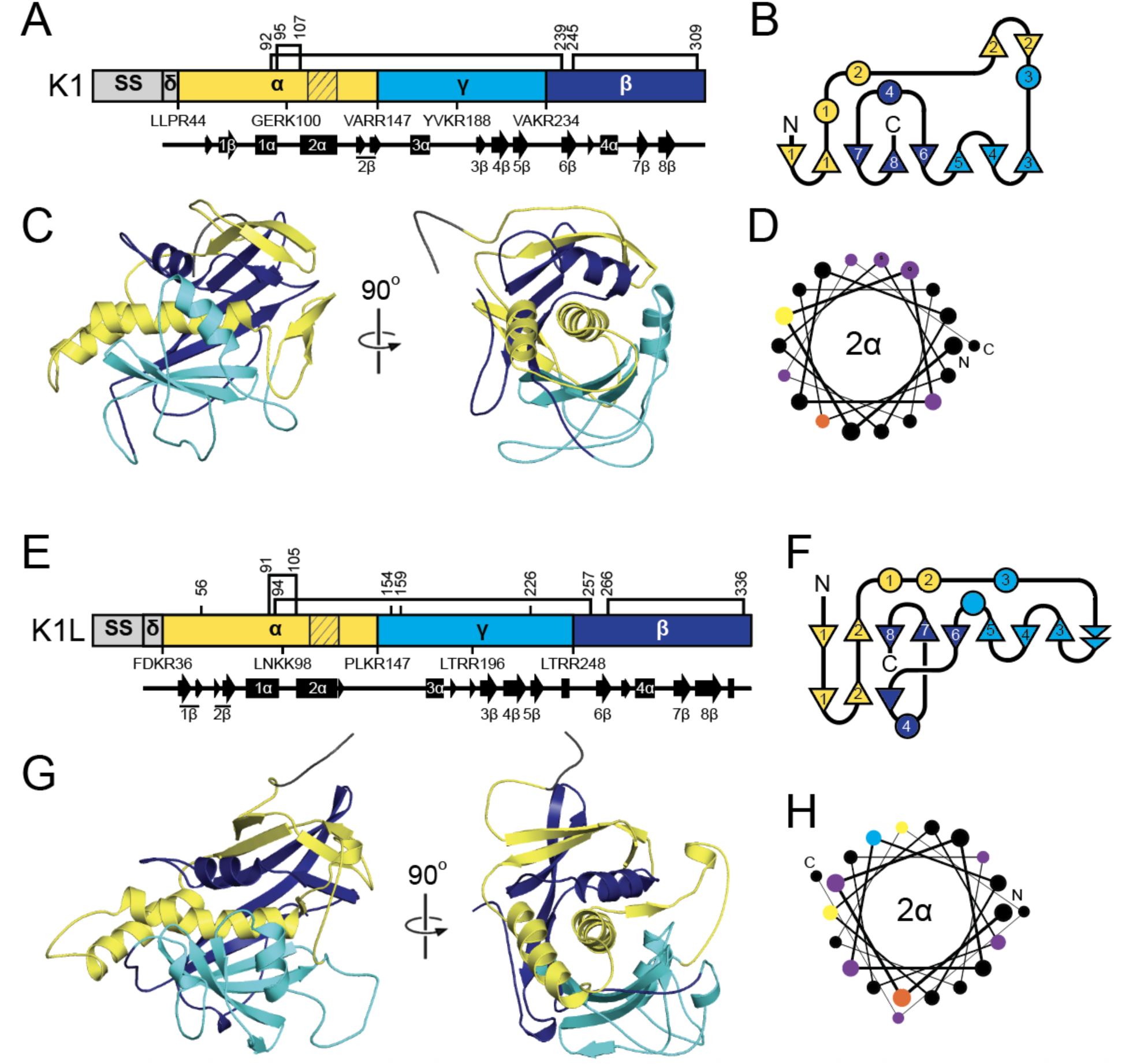
Secondary and tertiary structure models of K1 family killer toxins. (A, E) Domain diagrams of K1 and K1L ppTox indicating the sites of proteolytic processing that define the delta, alpha, gamma, and beta domains. The positions of Kex protease cleavage sites are indicated below the domain diagram, with the four amino acids preceding the cleavage point illustrated. Cysteine residues are indicated by lines and numbers above the domain diagram, with connections between them representing predicted disulfide bonds. Hatching in the diagram represents an amino acid sequence predicted to form transmembrane or pore-forming structures. Secondary structure, with numbered alpha helices and beta strands, is represented by arrows and rectangles below the domain diagram, respectively. (B, F) Two-dimensional schematic of the relative secondary structure organization of K1 and K1L with helices represented as circles and beta strands as triangles. N, amino-terminus, C, carboxyl-terminus. (C, G) Tertiary structure model of killer toxin pTox colored by domains as depicted in panels A and E. (D, H) Helical wheel diagram of 2α helix, with hydrophobic (black), positively charged (yellow), negatively charged (blue), polar (purple), and aromatic (orange) amino acids. The line thickness between amino acids (from thick to thin) represents the progression of the sequence from the N-terminus to the C-terminus.

As expected, K1L and its homologs have a similar secondary structure organization to K1, including delta, alpha, gamma, and beta domains and a similar pattern of six cysteine residues split evenly between the alpha and beta domains (Figure 2A). These domains were defined by putative Kex cleavage sites which resulted in similar sized domains to K1 [23,69]. K1L has longer delta, alpha, gamma, and beta domains (by 2, 6, 16, and 10 amino acids, respectively) compared to K1. Kex1 and Kex2 are also both required for the production of active K1L homologs [69] . The predicted alpha domain of K1L and its homologs are also cytotoxic when expressed by *S. cerevisiae*, which is consistent with the function of the K1 alpha as the toxin domain [51,69] . Overall, these empirical data support the close functional and structural relationship between K1, K1L, and their homologs.

#### 2.3.2 The K1 killer toxin family: Antifungal activity and immunity

The K1 alpha and beta heterodimer is an ionophoric toxin that increases the permeability of the yeast plasma membrane, resulting in cell death [70,71]. The mechanism of intoxication of K1 begins with binding and accumulation of the toxin at the cell wall [72]. Both domains of mature K1 (alpha and beta) have been implicated in cell wall binding. Fractionation of the yeast cell wall by chemical and enzyme extraction and competition assays with purified polysaccharides has identified that 1,6-beta-D-glucan is the primary receptor for K1 [73,74]. Moreover, manipulation of 1,6-beta-D-glucan content in the cell wall can alter susceptibility to K1, with depletion and overexpression leading to decreased or increased sensitivity to K1 intoxication, respectively. After cell wall binding, K1 attacks the plasma membrane, which is dependent on the Kre1 glycosylphosphatidylinositol (GPI) anchored protein [75,76]. Kre1 is thought to be the membrane receptor for K1, and deletion of the gene confers high levels of K1 resistance to both cells and spheroplasts (cells lacking a cell wall) [72,75–78]. A region of the C-terminus of Kre1 can bind K1 *in vitro* and is both necessary and sufficient for the intoxication of spheroplasts, but is insufficient for the intoxication of whole cells [76]. After Kre1 interaction, the exact mechanism of K1 intoxication is still unresolved.

The increased permeability of the cell membrane after K1 intoxication is thought to be due to the formation of voltage-independent ion channels that cause the energy-independent efflux of potassium ions and, potentially, the influx of hydrogen ions [79,80]. An alternative hypothesis for the observed ion leakage during K1 intoxication is the activation of the Tok1 potassium channel [81]. However, the role of Tok1 in K1 sensitivity has been questioned due to a lack of reproducibility in data showing the K1-resistance of *tok1Δ* strains [14,82]. Furthermore, *kre1Δ* spheroplasts are K1-resistant, even in the presence of *TOK1* [76]. Regardless of the exact mechanism of intoxication, the disruption of critical electrochemical gradients results in cell death by different pathways, depending on the K1 concentration [83,84]. The K1 alpha domain appears to be solely responsible for intoxication, as overexpression of this domain alone causes a suicidal phenotype in *S. cerevisiae* that is independent of the Kre1 membrane receptor [51]. The primary structure of K1 alpha has been noted to contain alpha helical regions of hydrophobicity that support a mechanism of membrane attack by K1. Some evidence for K1 oligomerization, consistent with channel formation, comes from observing K1 assembling into large complexes or aggregates of greater than 200 kDa, as well as soluble octamers [48,85,86].

The expression of ppTox K1 is necessary and sufficient for the self-protection of K1 killer yeasts against attack from exogenous K1 killer toxin. K1 ppTox immunity depends on export to the ER, but the signal sequence is not directly involved in immunity [87]. Early studies were able to decouple immunity from K1 toxicity by isolating non-toxic K1 mutants that provide immunity, including mutants with specific defects in pTox proteolytic processing. Similarly, yeast strains lacking functional *KEX1* or *KEX2* do not produce mature K1 but are immune to exogenous K1. Mutation of the beta domain or its deletion is also dispensable for K1 immunity, as is the majority of the gamma domain [87]. Moreover, truncation of K1 from the C-terminus identified that the minimal immunity domain of K1 consists of delta and alpha with 31 amino acids of gamma [67]. Mutagenesis further narrowed the minimal immunity region to the latter half of alpha and the N-terminus of gamma, and identified that amino acids C95 and C107 are essential for toxicity and immunity [87]. The attachment of lysozyme, random amino acids, or the gamma domain from K28 to the C-terminus of K1 alpha also maintained immunity [67,88] . Current working models of immunity predict a partially processed pTox (with alpha and gamma linked), preventing K1 intoxication at the cell surface by sequestering the Kre1 membrane receptor or mature K1 [67,87]. These models are supported by the detection of pTox mutants that provide K1 immunity outside of the cell; however, the exact mechanism of K1 immunity remains to be determined. However, the lethality of the K1 alpha domain has allowed the investigation of immunity mechanisms. Unlike exogenous K1, cell killing caused by endogenous K1 alpha appears to be independent of the Kre1 membrane receptor [51]. The lethality of K1 alpha can be suppressed by the co-expression of ppTox, but is dependent on cysteine residues C92, C95, and C107 in K1 alpha [51]. This observed “immunity” is also independent of Kre1, indicating that there could be a shared pathway that enables ppTox to protect cells from both endogenous K1 alpha and exogenous mature K1 toxin [88].

#### 2.3.3 The K1 killer toxin family: Molecular modeling

The tertiary structure models of K1 and K1L revealed a shared architecture centered on a globular one-layer alpha/beta sandwich fold that encapsulates the central 2α helix. In K1, 2α is predicted to participate in membrane interaction and pore formation, which is consistent with previously mapped hydrophobic regions of the alpha domain [66,87,89]. In both the K1 and K1L models, 2α is buried in a pocket formed primarily by antiparallel beta sheets, with support from alpha helices from the surrounding alpha, gamma, and beta domains. In K1, 2α is nested in a pocket of approximately 1698.0 Å^2^, shaped by three discontinuous beta strands and sheets (1β, 3–6β, and 7-8β), burying 83.8% of the helix (Table S6). In K1L, 2α is enclosed by a similar assembly of antiparallel beta strands from the alpha (1-2β), gamma (3-5β), and beta domains (6–8β) that buried 90.9% of the surface area of the helix (2186.8 Å^2^) (Table S6). The K1L alpha domain also formed a continuous beta sheet with strands 1-2β, in contrast to the discontinuous configuration of the analogous beta strands in K1.

The K1 and K1L models predicted both interdomain and intradomain disulfide bonds. In K1, a predicted disulfide bond between C92 (alpha domain) and C239 (beta domain) likely forms before Kex cleavage in the Golgi, stabilizing the heterodimer [62]. This prediction aligns with empirical data indicating that C239 is the only beta-domain cysteine essential for the alpha-beta disulfide linkage. The K1L model mirrors this architecture with a predicted disulfide between C94 and C257. Intradomain disulfides are also predicted in both toxins, such as C95–C107 in the alpha domain and C248–C312 in the beta domain of K1, which contribute to structural rigidity and functional properties such as immunity and cell binding [62].

Both structural models are consistent with the known posttranslational maturation pathway of K1-family toxins, in which the pTox is processed in the Golgi by Kex1 and Kex2 proteases. These cleavage sites occur at dibasic sites on solvent-exposed loops between domains, regions that are clearly accessible in tertiary structure models of K1 and K1L. Additional Kex cleavage sites are also located in the alpha and gamma domains on exposed linkers between 1α and 2α, and 3α and 3β. The internal cleavage of gamma appears to be important for K1 toxicity, but the functional relevance of the additional processing of alpha is unclear. A similar arrangement is observed in K1L, where the longest surface loops align with predicted Kex cleavage sites that define domain boundaries.

There are 22 point mutations from previous studies that reduce the toxicity, immunity, and cell wall binding properties of K1. To determine whether these mutations caused functional defects due to loss of protein stability, FoldX was used to predict the changes in folding stability (ΔΔG_fold_) of the K1 tertiary structure model (Table S4) [90]. The majority (15 of 22) were predicted to be destabilizing using a cutoff of ΔΔG_fold_ >2 kcal mol^-1^, and only four are not surface exposed (V116, S124, I151, C248). Most destabilizing mutations reduced toxin secretion (11/15) and antifungal activity against whole cells (15/15). Only two mutations predicted to be destabilizing (G264L and T191P) retained >75% of wild-type activity and secretion.

All but one of the seven single cysteine mutants in K1 are predicted to be structurally unstable, which is consistent with their general loss-of-function. All cysteine mutations cause a loss of K1 toxicity, and only C92S and C239S retained functional immunity. Empirical data show that cysteine mutations in the alpha domain (C92S, C92Y, C95S, and C107S) and beta domain (C239S) all reduce the expression of extracellular pTox and/or mature K1 [62,67]. The instability of alpha domain mutants likely explains the loss of K1 alpha toxicity when expressed alone in *S. cerevisiae*. Conversely, K1 with single (C248S) or double (C248S and C312S) mutations in the beta domain are expressed at wild-type levels and capable of inhibiting the growth of spheroplasts but not whole cells. This indicates that the predicted misfolding of the K1 beta domain would generally prevent cell wall binding but not cell membrane receptor binding. Therefore, these mutations appear not to cause additional misfolding of the alpha domain or loss of K1 expression, enabling membrane receptor binding and membrane permeabilization by the alpha domain.

Seven of the eight K1 mutations predicted to be stable ((ΔΔG_fold_ <±2 kcal mol^-1^) resulted in defects in toxicity and/or immunity (V85T, D101R, S124P, I129R, D140R, N181K, and R188A) (Table S4). Six of these amino acids were predicted to be surface-exposed (V85T, D101R, I129R, D140R, N181K, and R188A), and three were positioned in regions of the protein lacking secondary structure (I129R, N181K, R188A). These mutations could therefore be important for defining areas of K1 that are important for cell recognition or for conformational changes required for toxicity (Figure 3). Specifically, residue D101 in the alpha domain is surface-exposed and positioned on the loop between helices 1α and 2α. Notably, the mutation D101R (ΔΔG_fold_ = 0.52 kcal mol^-1^) results in a loss of cell wall binding while retaining the ability to kill spheroplasts and confer immunity. This residue could define a surface contact point between the toxin and the yeast cell wall that could be useful in defining K1 specificity.

**Figure 3.**
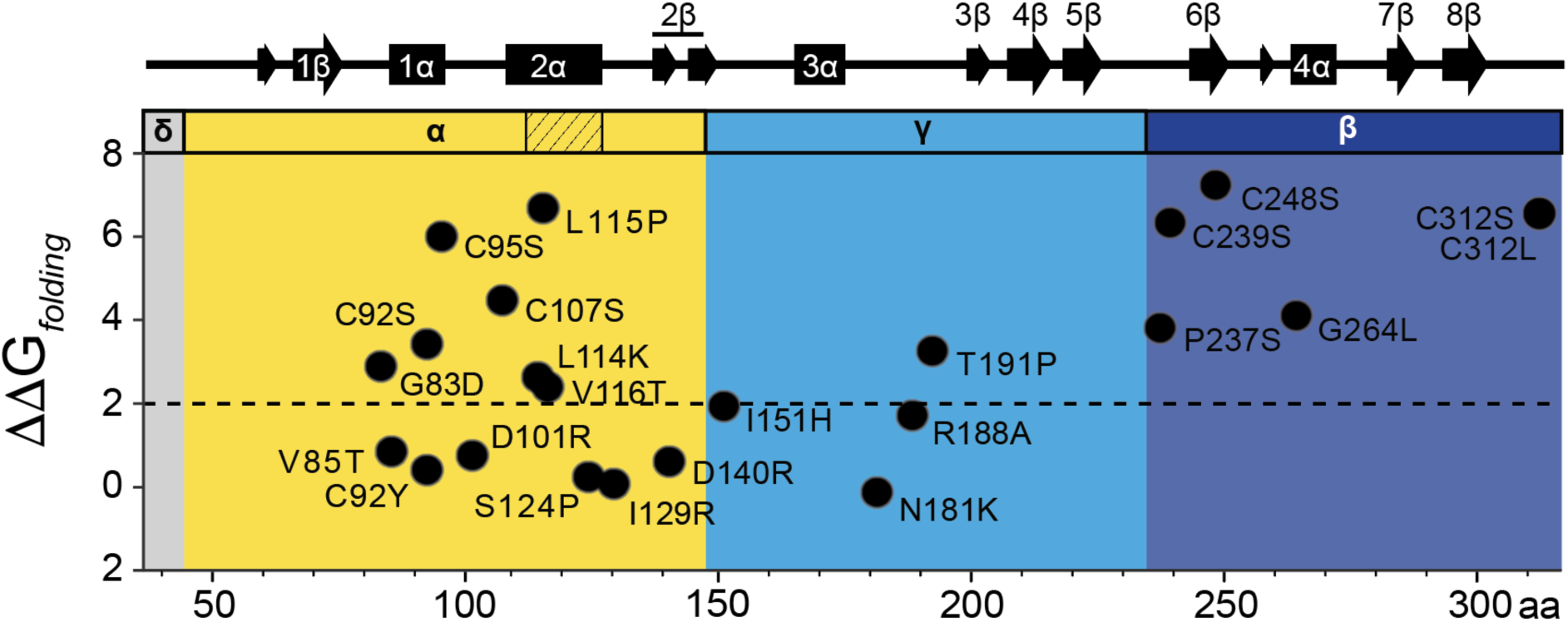
FoldX analysis of K1 mutations identified in previous empirical studies. (A) FoldX mutagenesis of published point mutations positioned relative to the alpha, gamma, and beta domains and secondary structure of K1. The cutoff for mutations that are predicted to disrupt protein structure is shown as a red dotted line (Λ1Λ1G_fold_ ±2 kcal mol^-1^). K1 secondary structure with alpha helices and beta strands are shown as rectangles and arrows above each domain, respectively.

### 2.4 The K2 killer toxin family

K2 was the second killer toxin discovered after a screen of 964 yeasts of various genera from the National Collection of Yeast Cultures (NCYC). All K2 killer yeast strains originated from ale yeasts isolated from U.K. breweries, and K2 was identified as a toxin with a distinct activity and immunity profile [91]. Since their initial discovery, K2 killer yeasts have been repeatedly isolated from wineries and breweries [37,92–94]. Extraction of dsRNA from K2 killer yeasts confirmed the presence of an M2 dsRNA satellite. The dsRNA was required for K2 expression, as curing with cycloheximide or high temperatures caused the loss of M2 and the K2 killer phenotype [1]. The genetic sequence of the M2 satellite identified the K2 open reading frame that was later confirmed to encode the K2 ppTox responsible for the antifungal and immunity phenotypes [95–97]. K2 displays optimal killing activity at pH 4.3 and 20-25°C [50]. It has a broad spectrum of activity, inhibiting the growth of many killer and non-killer strains of species in the *Saccharomyces* genus as well as the human pathogen *N. glabratus* and diastatic brewing strains [28,91,98].

The K21-producing strain *S. paradoxus* T21.4 was originally isolated from oak trees in the U.K. [99]. The toxin produced by *the S. paradoxus* T21.4 was erroneously considered a K1 or K28 toxin [100,101] before being designated the unique toxin named K21 [100]. K21 was confirmed to be encoded by an M satellite by dsRNA extraction and curing of the dsRNA satellite by growth at elevated temperature [101]. Sequencing of the M21 satellite showed little nucleotide sequence homology to known yeast killer toxins but had a similar organization to other dsRNA satellites [100]. Another dsRNA-encoded toxin, K66, with 92% amino acid identity to K21, was later identified in the *S. paradoxus* strain ALM–66 from the spontaneous fermentation of serviceberries [16]. The antifungal activities of K21 and K66 inhibit the same species of yeasts, and the immunity functions of both toxins are cross-protective (i.e., K21 protects against K21 and K66). K66 exhibits optimal antifungal activity at 20°C at pH 4.8 [16] .

First discovered in 1990, KHS (killer of heat-sensitive) was identified in *S. cerevisiae* isolated from Japanese wineries [102]. The killer yeast was found to have no M satellite, and the killer phenotype was resistant to curing by cycloheximide treatment and incubation at elevated temperatures, suggesting it was a genome-encoded killer toxin [102]. The *KHS1* gene was initially identified from *S. cerevisiae* genomic libraries and mapped to the right arm of chromosome V [103]. This gene was cloned and expressed ectopically, confirming that it conferred the killer phenotype and its respective immunity functions [103]. Initial sequencing data were found to contain errors, which were later corrected in subsequent studies [41] . Importantly, *KHS1* is absent from the reference genome of *S. cerevisiae*, but is present in most other strains, often with polymorphisms and premature stop codons [104,105]. The close similarity of a subset of *KHS1* genes identified in *S. cerevisiae* and *S. paradoxus* supports introgression from *S. paradoxus* [104] . Homologs of *KHS1* have also been identified in the genomes of many different yeast species within the Saccharomycotina [41] .

The antifungal activities of KHS are consistent with other killer toxins, with an optimal activity at pH of 4.7 and ambient temperatures <30°C. Comprehensive screening of more than 1,000 strains of *S. cerevisiae* confirmed the prevalence of killer toxin production that correlated with functional *KHS1* [37,104]. However, the antifungal activities of KHS appear to be ineffective against many strains of *S. cerevisiae* and *S. paradoxus*, likely due to the prevalence of *KHS1* and any associated immunity function. Thus, prior screens for killer toxins have likely failed to recognize the widespread production of active KHS by *S. cerevisiae*. Indeed, the antifungal activities of KHS so far appear limited to the opportunistic pathogen *N. glabratus* and a single strain of *S. cerevisiae* [37,103]. Yet little is known about the mechanism of action of KHS, although its amino acid sequence homology to K2 and K21 suggests a similar mechanism of action.

#### 2.4.1 The K2 killer toxin family: Domain organization and posttranslational modification

The K2 ppTox consists of 362 amino acids across four domains: delta (amino acids 1-79), alpha (80-165), gamma (166-268), and beta (269-362), in order from the N- to C-terminus (Figure 6A). The primary transcription product is predicted to encode a 39 kDa protein [96]. Similar to K1, the K2 ppTox enters the secretory pathway, first being exported to the ER, where a signal sequence of undetermined length is cleaved. The signal sequence prediction tools SignalP and PSIPRED fail to recognize a canonical signal sequence in K2; however, the N-terminal prepro region possesses properties consistent with an *S. cerevisiae* signal sequence [106]. These include an N-terminal region with positively charged residues, followed by a hydrophobic alpha-helical region, and a recognition sequence for a signal peptidase. The difficulty in detecting this signal sequence in K2 is due to an additional ∼30 residues before the predicted cleavage site, rather than the typical five residues. *In silico* truncation of the K2 N-terminus results in the recognition of a signal sequence by prediction tools [106]. This analysis predicted that the signal peptidase cleavage site for K2 is between or after amino acids 52-56.

**Figure 6.**
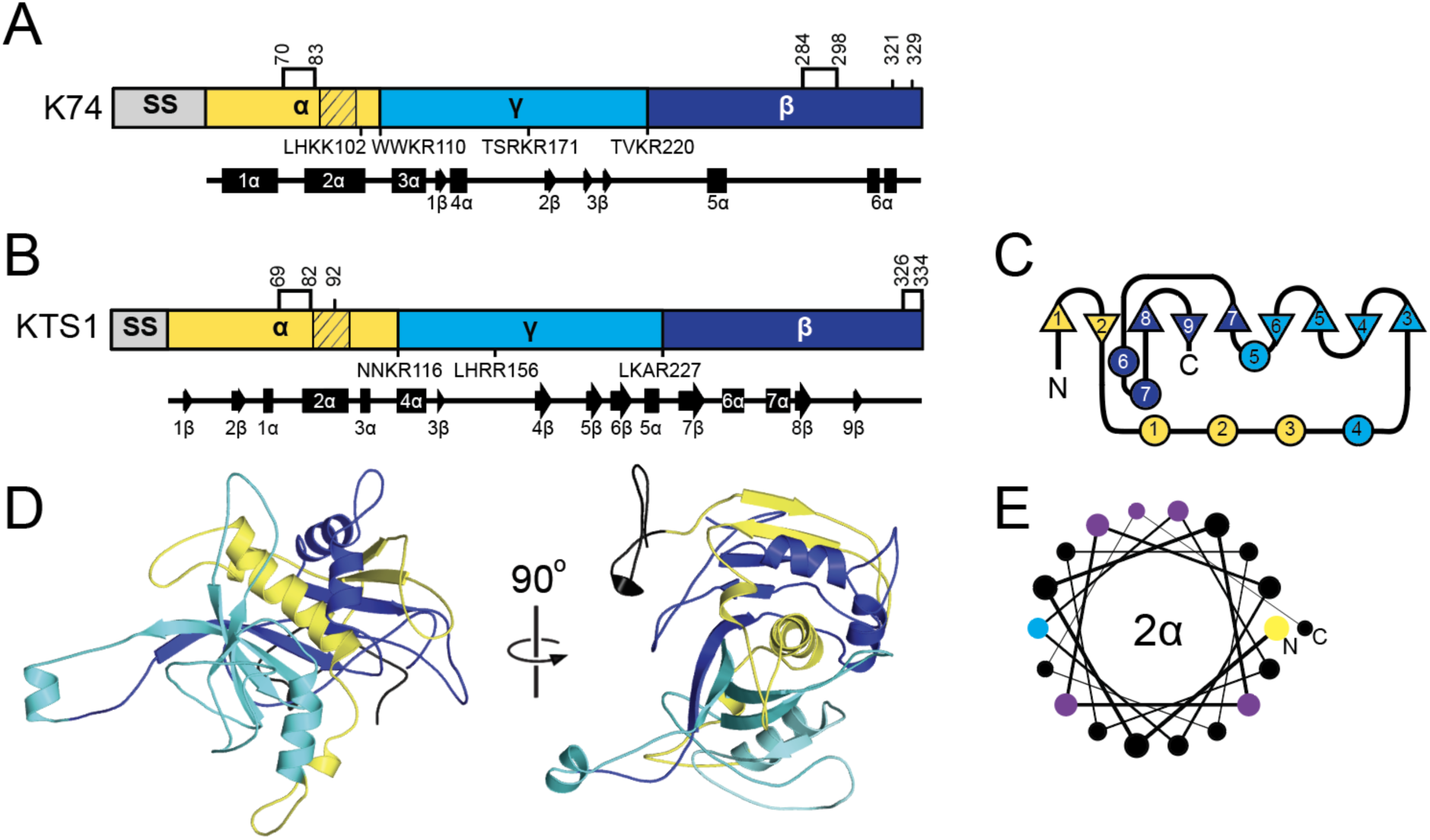
-Secondary and tertiary structure models of K74 family killer toxins. Domain diagrams of (A) K74 and (B) KTS1^Cmal^ ppTox indicating the sites of proteolytic processing that define the delta, alpha, gamma, and beta domains. The positioning of Kex protease cleavage sites is indicated below the domain diagram, with the four amino acids preceding the cleavage point illustrated. Cysteine residues are indicated by lines and numbers above the domain diagram, with connections between them representing predicted disulfide bonds. Hatching in the diagram represents amino acid sequences predicted to form transmembrane or pore-forming structures. Secondary structure, represented by numbered alpha helices and beta strands, is illustrated by arrows and rectangles below the domain diagram, respectively. (B) Two-dimensional schematic of the relative secondary structure organization of KTS1^Cmal^ with helices represented as circles and beta strands as triangles. N, amino-terminus domain, C, carboxyl-terminus. (C) Tertiary structure model of killer toxin pTox colored by domains as depicted in panels A. (D) Helical wheel diagram of 2α helix, with hydrophobic (black), positively charged (yellow), negatively charged (blue), and polar (purple) amino acids. The line thickness between amino acids (from thick to thin) represents the progression of the sequence from the N-terminus to the C-terminus.

Following K2 signal peptidase cleavage, pTox enters the ER, where disulfide linkages can be formed in a manner similar to other killer toxins. K2 pTox in the Golgi would be predicted to be cleaved by the Kex2 protease after R79, R165, R221, and R268, with likely additional processing by Kex1, resulting in a mature heterodimeric toxin [96]. Loss of either or both Kex proteases reduces or eliminates K2 toxicity, respectively, without altering K2 immunity function [96]. Assuming processing at the non-canonical signal peptidase cleavage site and predicted Kex cleavage sites, K2 alpha and beta have theoretical molecular weights of ∼8.7 kDa and ∼10.5 kDa, respectively. Mature K2 appears to have an apparent molecular weight of ∼21.5 kDa. Theoretical predictions do not agree well with a previously tagged C-terminal beta domain with an apparent molecular weight of ∼17 kDa, which would be more consistent with a beta domain attached to a partially cleaved or uncleaved gamma domain (15.8 and 21.6 kDa, respectively). However, an extensive mutagenesis study supports a K2 domain organization similar to that of K1/K28, with a gamma domain that is internally cleaved and removed during the maturation of the toxin [107] . This is evident in the gamma domain’s high mutational tolerance and the observed decrease in functionality resulting from the mutation of Kex cleavage sites [107]. However, the amino acid sequence of the terminal ends of the mature K2 toxin remains to be experimentally validated.

The K21 ppTox consists of a 346 amino acid protein that can be divided into four domains; delta (amino acids 1-59), alpha (60-129), gamma (130-239), and beta (240-346) in order from the N-to C-terminus (Figure 7A). Domain boundaries are defined by dibasic and basic residue motifs for Kex cleavage that are positioned similarly to K2. PSIPRED predicts the K21 signal sequence to be after amino acid 40, in line with the organization and posttranslational modification of K2. Moreover, K21 has a hydrophobic region between residues 23-39 that is typical of a signal sequence, but like K2, is positioned away from the N-terminus. K21 also has a region of positive charge with amino acids R9, R14, and K22 forming a conserved triad of residues before the hydrophobic region in the N-terminus. Although K21 has only four cysteine residues, their positioning in the alpha and beta domains is similar to K2.

**Figure 7.**
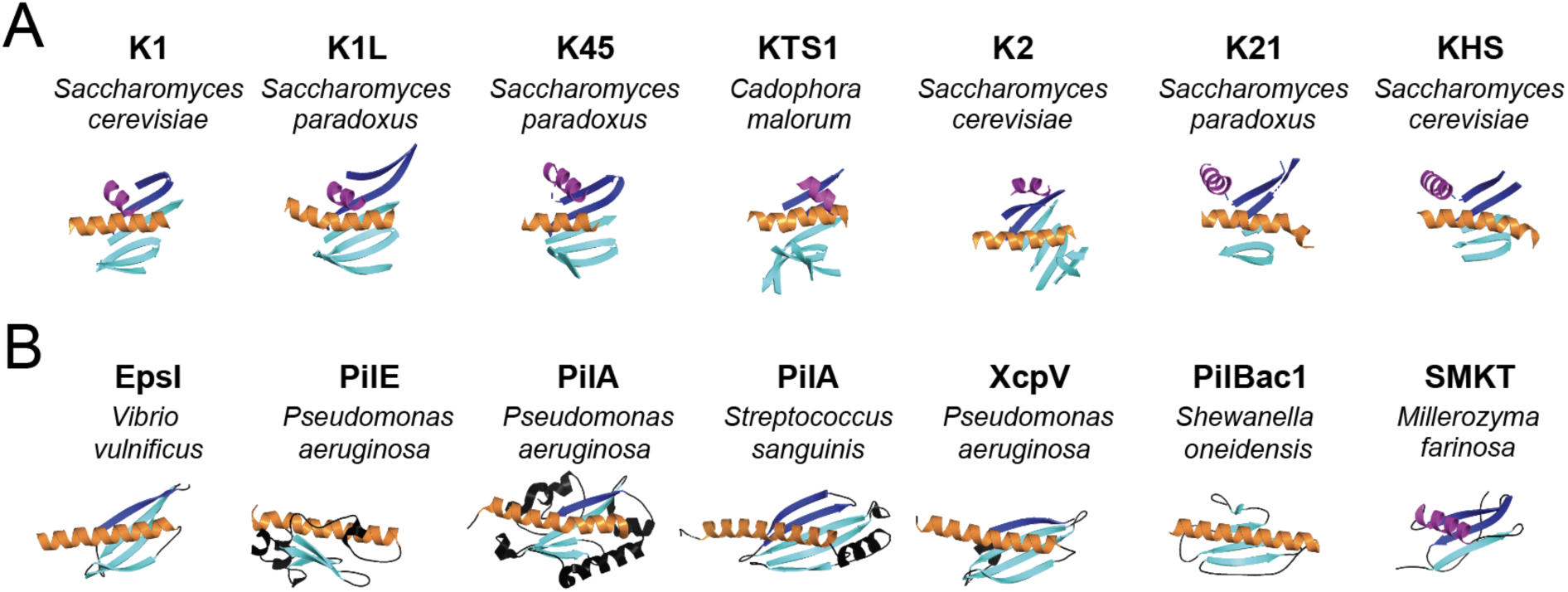
A conserved core motif of the K1 superfamily shows structural similarity to pilins, pseudopilins and a killer toxin from *Millerozyma farinosa*. (A) Core K1 superfamily toxin motifs and (B) bacterial pilins, pseudopilins from *Pseudomonas aeruginosa* (PilE (PDB: 4noa), PilA (PDB: 3jyz), XcpV (PDB: 5bw0), *Streptococcus sanguinis* (PilA (PDB: 7o5y)), *Shewanella oneidensis* (PilBac1 (PDB: 4d40)), *Vibrio vulnificus* (EpsI (PDB: 2ret)) and a killer toxin from *Millerozyma farinosa* (SMKT (PDB: 1kvd)) identified by DALI analysis as homologs of K1 superfamily killer toxins. Secondary structure elements that are conserved across structures are colored to highlight similarities including the central hydrophobic helix (orange) and a second conserved helix (magenta) that are positioned before and after beta strands (cyan) ending in C-terminal beta strands (dark blue).

There has been no prior attempt to determine whether KHS is posttranslationally modified, as are other canonical killer toxins. However, due to its sequence homology with K2 and K21, it is predicted to have four domains: delta (1-63), alpha (64-130), gamma (131-237), and beta (238-350). Unlike K2 and K21, all of the domain boundaries are defined by dibasic residue motifs that are likely cleavage sites for Kex proteases. The positioning of the KHS dibasic sites supports the functionality of non-canonical monobasic cleavage sites of K2 and K21. Similar to K21, the signal sequence cleavage site of KHS is predicted to be positioned away from the N-terminus due to the extension of the N-terminus before the canonical pattern of positively charged residues (R20, R23, R26) and a hydrophobic domain (residues 27-46). PSIPRED predicts this signal peptidase cleavage site to be after residue 36. The pattern of cysteine residues in KHS is more similar to K21 than K2.

#### 2.4.2 The K2 killer toxin family: Antifungal activity and immunity

Similar to K1, the mature K2 toxin is thought to be an ionophore. The mature heterodimeric toxin interacts with 1,6-beta-D-glucan in the yeast cell wall as a primary receptor [108]. Enzymatic removal of the cell wall does not protect from K2 intoxication, indicating that K2 also interacts with the plasma membrane. Moreover, the loss of the Kre1 provides protection against K2 for whole cells and spheroplasts, and it has been suggested that, like K1, Kre1 is the secondary membrane receptor for K2 [109]. However, unlike K1, there has not been a direct measure of K2 interaction with Kre1. Exposure of yeast cells to K2 leads to cellular damage (as measured by lipophilic anion binding), which correlates to reduced respiration activity and lowered intracellular ATP levels, but without detectable ATP leakage seen during K1 intoxication [110]. Scanning electron microscopy of K2-intoxicated cells reveals shrinkage, loss of turgor, and cracks and pores on the cell surface, suggesting disruption of the cell wall and membrane, possibly induced by pore formation and changes in membrane permeability [93,111]. Transmission electron microscopy also revealed disruption of the cell wall and an abnormal undulating morphology of the plasma membrane. Expression of the K2 alpha domain is also toxic to yeasts, suggesting that it is responsible for the cytotoxicity of K2, as is the case for the K1 alpha domain [106] .

Genome-wide screens have identified hundreds of genes that influence K1, K2, and K21/K66 resistance or hypersensitivity [14,16,17]. Genes associated with resistance predominantly involve cell wall and plasma membrane structure, biogenesis, and mitochondrial function. Conversely, genes associated with hypersensitivity are primarily linked to stress signaling pathways and ion and pH homeostasis. These findings suggest that, while K2 and K21 share some mechanistic similarities with K1, supporting their classification as ionophoric toxins, they have unique interactions with specific cellular components during cellular intoxication.

Like the majority of killer toxins, K2 killer yeasts are immune to their own toxin [1]. Unlike K1 and K28 that require the ppTox for functional immunity, the prepro region at the N-terminus of K2 is necessary and sufficient for immunity [96,106] . The prepro immunity function of K2 was first observed after the creation of a mutant that lacked K2 immunity but expressed an active toxin [96]. Moreover, yeast expressing this mutant K2 toxin grew well at pH 7 (K2 inactive), but were unhealthy at pH 4.5 (K2 active). A more recent study identified an N-terminal immunity peptide in the K2 prepro region, released after signal sequence cleavage [106]. The mechanism by which the K2 prepro region functions as an immunity factor is not fully understood; however, due to its hydrophobicity, it may interact with cell membranes, similar to other small immunity peptides from bacteriocin immunity systems [112]. Localization of the K2 immunity domain to membranes could prevent pore formation, but other mechanisms are also plausible. Despite the low sequence similarity and identity of 31% and 10%, respectively, K21 and KHS have a similar organization in the N-terminal region, suggesting a similar immunity mechanism to K2.

Toxins of the K2 family may share further functional similarity due to the presence of a conserved Pfam Domain of Unknown Function (DUF5341) (Figure 4A) [100]. This domain resides in the C-terminus of each K2 family toxin, beginning in the gamma domain and extending nearly the entire length of the beta domain. This configuration predicts that complete cleavage by Kex proteases of the K2 family of toxins would split the domain into two, with only the C-terminal portion present in the mature toxin. DUF5341 has 106 Uniprot entries, 82 of which are uncharacterized fungal proteins with unknown function in the genomes of the Ascomycota <u>Table SX</u>. The remaining 24 entries include other members of the K2 family, proteins that appear to be associated with the fungal cell wall, and several YER187W and YGL262-like proteins, whose similarity to KHS and K2 has been previously documented [41]. The majority (76%) of DUF5341 domains are found in the C-terminus of small proteins (100-400 amino acids), similar to the organization found in the K2 family. Exactly half of the DUF5341-containing proteins also overlap with PSI-BLAST hits for the K2 family and are mostly uncharacterized. While the function of DUF5341 is unknown, homology to the beta domain of the K2 family may suggest a conserved carbohydrate-binding functionality across these proteins.

**Figure 4.**
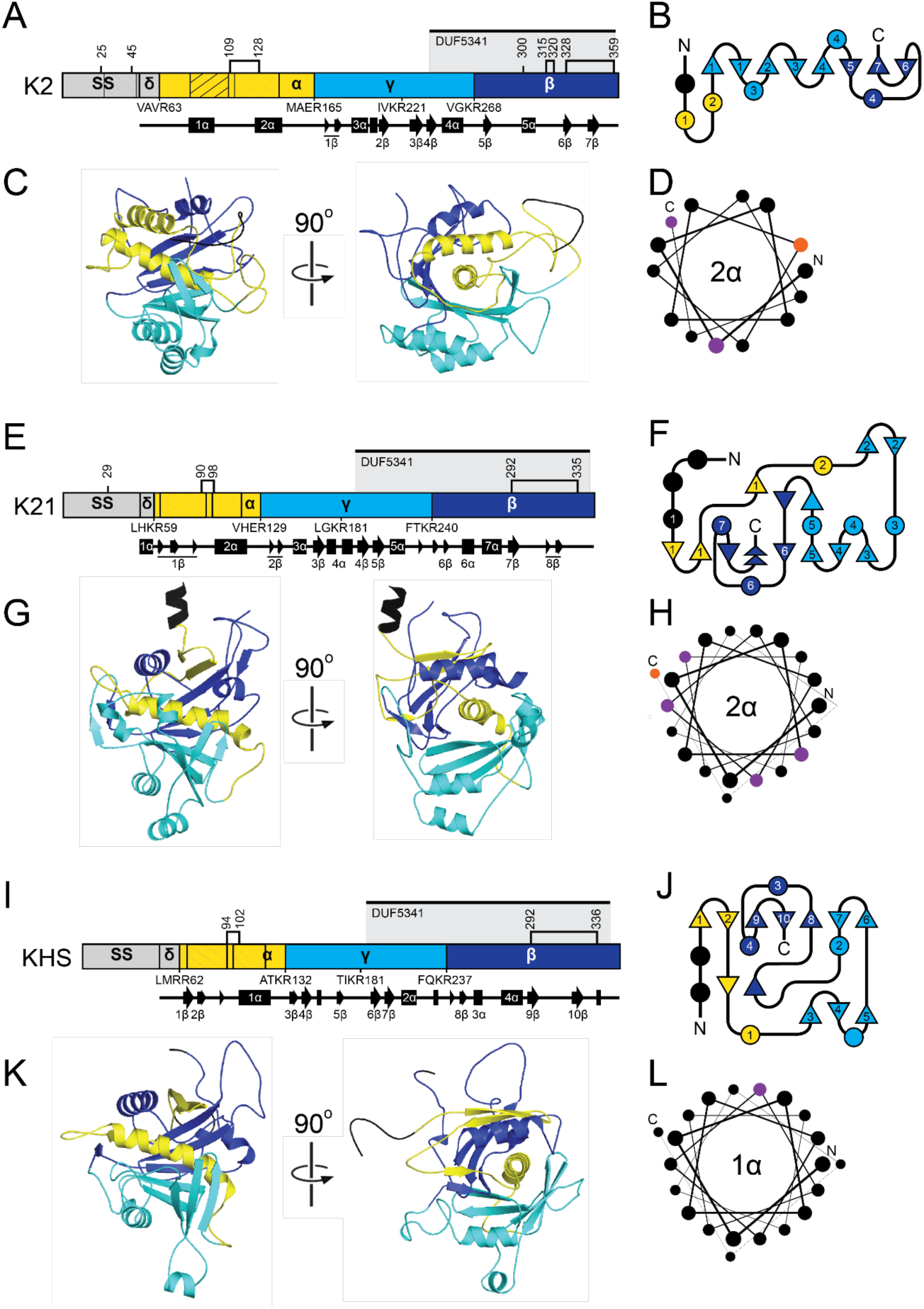
-Secondary and tertiary structure models of K2 family killer toxins. (A, E, I) Domain diagrams of K2, K21, and KHS ppTox indicating sites of proteolytic processing that define the delta, alpha, gamma, and beta domains. The positioning of Kex protease cleavage sites is indicated below the domain diagram, with the four amino acids preceding the cleavage point illustrated. Cysteine residues are indicated by lines and numbers above the domain diagram, with connections between them representing predicted disulfide bonds. Hatching in the diagram represents amino acid sequences predicted to form transmembrane or pore-forming structures. Secondary structures, including numbered alpha helices and beta strands, are represented by arrows and rectangles, respectively, below the domain diagram. DUF5341 aligning region marked by a bar above the domain diagrams at amino acids 239-360 (K2), 190-343 (K21), and 185-344 (KHS). (B, F, J) Two-dimensional schematic of the relative secondary structure organization of K2, K21, and KHS, with helices represented as circles and beta strands as triangles. N, amino-terminus domain; C, carboxyl-terminus. (C, G, K) Tertiary structure models of K2, K21, and KHS pTox colored by domains as depicted in panels A, E, and I. (D, H, L) Helical wheel diagram of 2α helix, with hydrophobic (black), polar (purple), and aromatic (orange) amino acids. The line thickness between amino acids (from thick to thin) represents the progression of the sequence from the N-terminus to the C-terminus.

#### 2.4.3 The K2 killer toxin family: Molecular modeling

Superimposition of the predicted structures of the K2 family resulted in RMSD values of 5.8 Å (K2/K21), 3.9 Å (K2/KHS), and 1.1 Å (K21/KHS) and revealed a shared globular organization centered around a conserved central alpha helix, which is surrounded asymmetrically by a series of discontinuous antiparallel beta-strands. This one-layer alpha/beta sandwich architecture appears characteristic of *Saccharomyces* ionophoric toxins and is consistent with the predicted structures of the K1 killer toxin family.

In K2, the central 2α helix is positioned within the alpha domain and is flanked by a total of eight beta strands: 1-2β (alpha domain), 3-4β (gamma domain), and 5-7β (beta domain). Helices from all three domains, including 1α, 3α, and 4α, contribute to wrapping the central helix and the burial of 1652.2 Å^2^ or 95.5% of the central helix (Table S6). Notably, helices 5α and 6α lie on the side opposite the beta sheet and do not appear to interact directly with 2α. Hydrophobicity and PSIPRED analyses suggest that helix 2α and 1α of the alpha domain may function as membrane-interacting or pore-forming structures, reinforcing the proposed mechanism of action for K2. The AlphaFold2 model of K2 is consistent with its known Golgi-mediated post-translational processing, as experimentally validated Kex cleavage sites are located on solvent-exposed loops that would facilitate proteolysis (Table S5). Unlike K1, K2 does not appear to form interdomain disulfide bonds between the alpha and beta domains. Instead, intradomain disulfide bonds are predicted in the alpha domain (C109–C128) that links the N-terminus of 2α to the C-terminus of 1α. One of the two disulfide bonds in the beta domain serves to link the K2 C-terminal tail to the penultimate beta strand (6β), reminiscent of K1 pTox. Recent studies support the functional significance of these cysteines in K2 activity [107].

The K21 structural model closely mirrors that of K2 but displays more disordered regions and less extensive secondary structure. Like K2, K21 has a central 2α helix (residues 97-119) predicted to be transmembrane based on hydrophobicity and secondary structure predictions. The central helix is embedded in a seven-stranded β-sheet composed of elements from all three domains: 1-3β (alpha domain), 4-5β (gamma domain), and 6-8β (beta domain) that bury a surface area of 1652.2 Å^2^ or 83.2% of the central helix (Table S6). Surrounding helices (5α, 6α, and 7α) interact with exposed regions of the 2α helix, while 3α and 6α interact with the solvent-exposed face of the gamma domain. An additional hydrophobic region with beta strands, located at residues 62-85 in the alpha domain, aligns with a similar helical region in K2, suggesting that it may contribute to membrane association. K21 is predicted to undergo Kex cleavage at exposed loops, and intradomain disulfide bonds are formed in the alpha and beta domains, similar to K2.

The KHS model shares the same core topology with K2 and K21, though with key differences. It features a central alpha-helix (1α) surrounded by nine antiparallel beta-strands: 1-2β (alpha domain), 6-7β (gamma domain), and 8-10β (beta domain), burying a large area of 1831.8 Å² or 89.2% of the surface area of 1α (Table S6). Like K2 and K21, the central helix of KHS is predicted to be transmembrane, being almost exclusively composed of hydrophobic residues (Figure 4L). A second transmembrane regions is also present in the alpha domain, consisting of beta sheets similar to K21 (amino acids 68-94). Helices 4α and 5α (from the alpha and beta domains) contact the exposed side of the central helix. Similar to K21, KHS shows increased structural disorder and loop flexibility compared to K2. Kex cleavage sites and disulfides are also positioned similarly to K21 and K2.

### 2.5 The K45 killer toxin family

K45 is a poorly studied killer toxin that is produced by *S. paradoxus* N-45, which was isolated from the exudate of a Mongolian oak (*Quercus mongolica*) in Ternei City, Russia [99] (see ARS Culture Collection (NRRL)). After an initial 2013 screening that failed to identify N-45 as a killer yeast, an antifungal phenotype was observed in a later survey, which mischaracterized the toxin as K1 [113]. A more detailed analysis of this killer yeast revealed a dsRNA satellite, designated M45, that showed little nucleotide homology to either M28 or M1, as determined by northern blotting [100]. Further efforts to obtain the genetic sequence of M45 identified the killer toxin K45 but no other studies have been undertaken to characterize this toxin.

#### 2.5.1 The K45 killer toxin family: Domain organization and posttranslational modification

Overall, the domain organization of K45 is predicted to be similar to other *Saccharomyces* killer toxins, with the order of delta, alpha, gamma, and beta. The boundaries between these domains are defined by putative Kex cleavage sites, located between alpha and gamma after residue 179 and between gamma and beta after residue 267 (Figure 5A). The dibasic motif, comprising residues K96 and R97, which are located near the center of the alpha domain, is similar to dibasic sites identified in the alpha domains of KHS and K1, but with no known biological function. Although SignalP fails to identify a signal sequence cleavage site, PSIPRED predicts cleavage after amino acid 27, and the preceding sequence exhibits an organization typical of a *Saccharomyces* signal sequence, featuring positively charged amino acids and a hydrophobic region. Similar to K1 and K2 toxin families, K45 contains a hydrophobic helix, which is in the C-terminal half of the alpha domain. A second hydrophobic region in the alpha domain encompasses the beta strand 2β.

**Figure 5.**
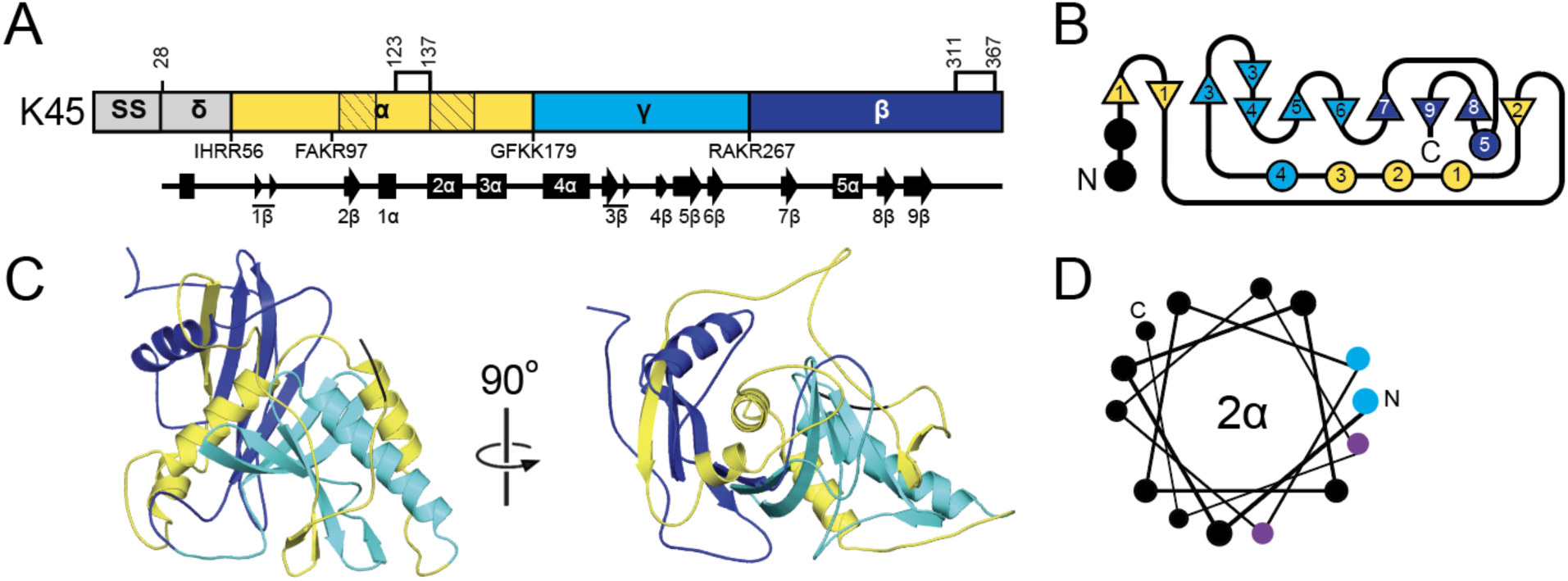
-Secondary and tertiary structure models of K45 family killer toxins. (A) Domain diagrams of K45 ppTox indicating the sites of proteolytic processing that define the delta, alpha, gamma, and beta domains. The positioning of Kex protease cleavage sites is numbered below the domain diagram with the four amino acids prior to the cleavage point illustrated. Cysteine residues are indicated by lines and numbers above the domain diagram, with connections between them representing predicted disulfide bonds. Hatching in the diagram represents an amino acid sequence predicted to form transmembrane or pore-forming structures. Secondary structure, with numbered alpha helices and beta strands, is represented by arrows and rectangles below the domain diagram, respectively. (B) Two-dimensional schematic of the relative secondary structure organization with helices represented as circles and beta strands as triangles. N, amino-terminus domain, C, carboxyl-terminus. (C) Tertiary structure model of killer toxin pTox colored by domains as depicted in panels A. (D) Helical wheel diagram of 2α helix, with hydrophobic (black), negatively charged (blue), and polar (purple) amino acids.The line thickness between amino acids (from thick to thin) represents the progression of the sequence from the N-terminus to the C-terminus.

#### 2.5.2 The K45 killer toxin family: Molecular modeling

K45 shares tertiary structure features with the K1 and K2 families, with a central alpha-helix (2α) surrounded by eight beta-strands organized in an antiparallel arrangement: 2β (alpha domain), 3-6β (gamma domain), and 7-9β (beta domain). This extensive structure, along with two other alpha helices (1α and 5α), buries 1158.1 Å² of surface area or 89.2 % of the surface area of the 2α helix (Table S6). The lone 5α helix of the beta domain is positioned antiparallel to the 2α helix and is offset by ∼20°. Compared to 2α, 5α is only partially wrapped by the central beta sheet, with a large portion of the structure being solvent-exposed. The hydrophobic 2β strand from the alpha domain is responsible for the majority of the interactions with 5α, as it is positioned at the end of the beta sheet region next to 8β. K45 has five cysteines, one of which is removed upon cleavage of the signal sequence, and the other four are predicted to form disulfide bonds in the alpha and beta domains. The organization of the cysteine and disulfide bonds is most similar to K2 family toxins. As with the K1 and K2 families, disulfides link the N-terminus of the central helix to the C-terminal end of the preceding secondary structure element (1α in K45) and pin the C-terminal tail to the beta domain.

### 2.6 The K74 killer toxin family

*S. paradoxus* Q74.4 and Y8.5 were both isolated from oak trees in the U.K. and identified as killer yeasts [101,113]. Despite the erroneous initial identification as K1, the novel spectrum of antifungal activity led to the discovery of K74 and the novel dsRNA satellite named M74 [100,114] . Efforts to study the function of K74 have been aided by the expression of the toxin by laboratory strains of *S. cerevisiae* and the cytoduction of M74. Overexpression of K74 determined that the optimal conditions for the activity of K74 are pH 4.3 and 20°C, which is consistent with other *Saccharomyces* killer toxins [114].

#### 2.6.1 The K74 killer toxin family: Domain organization and posttranslational modification

All analysis of K74 was reported by Rodriguez-Cousiño et al., which predicted posttranslational modification by proteolysis due to the presence of dibasic motifs [100,114]. These predictions would mean that K74 is similar in organization to a typical killer toxin with alpha, beta, and gamma domains, but with a significantly longer beta domain and shortened gamma domain when compared to other Saccharomyces killer toxins (Figure 6A). Expression of K74 in a kex2Δ null strain caused the loss of mature K74 and the accumulation of glycosylated pTox. Mutation of arginine residues (R110 and R220) at dibasic motifs that define the boundaries of alpha/gamma and gamma/beta domains, respectively, prevented the processing of the mature K74. Western blot analysis of K74 with reducing agents revealed the release of a ∼13 kDa C-terminal beta domain, suggesting that the mature K74 is a disulfide-linked heterodimer. Systematic mutation of all six cysteines of K74 revealed that they are all required for the production of the mature toxin; however, it remains unresolved whether any cysteine pair (or pairs) is responsible for intramolecular crosslinking between the alpha and beta domains. The positioning of the cysteines in the alpha and beta domains is similar to that of the K1/K2/K45 killer toxin families.

#### 2.6.2 The K74 killer toxin family: Molecular modeling

The confidence of the AlphaFold2 molecular model of K74 was the second lowest of all *Saccharomyces* killer toxins and had the highest proportion of unstructured loops (64.26%). (Table S3). AlphaFold3 was used to model K74, but also yielded a low-confidence model to create a more confident representative model of the K74 killer toxin family, 32 tertiary structure models were generated from K74 homologs (Figure 1). These sequence homologs had an average sequence identity of 19.2% (SD 14.8%). Molecular models were generated for all homologs, and 21 had high confidence scores with pTM values greater than 0.7 (Figure S3). These models contained putative Kex2 cleavage sites that divide the protein into a similar domain configuration as K1, featuring a central 2α helix from the alpha domain wrapped by beta strands of the alpha, beta, and gamma domains (Figure 6F). An uncharacterized K74 homolog from the saprophytic plant pathogen *Cadophora malorum* (Killer Toxin 74 from *C. malorum*; KTS1^Cmal^) was the model with the highest identity to K74, with a pTM score above 0.8. MD simulations indicated that the predicted structure was stable (Figure S2). All confident structures of K74 homologs have structural similarity to KTS1^Cmal^, with an average RMSD of 2.06 Å. Although these data indicate that K74 is an outlier in a larger family of K74-like proteins, the structures of homologous proteins provide a useful framework to further investigate the structure and function of K74.

The tertiary structure model of KTS1^Cmal^ is a globular protein with a similar organization to the ionophoric killer toxins (delta, alpha, gamma, and beta). The alpha and gamma domain boundary is a dibasic motif, whereas the boundary between gamma and beta is a non-canonical cleavage site (LKAR). Of the K74 homologs, 12 share the monobasic non-canonical gamma-beta cleavage site while 20 possess a dibasic cleavage site, suggesting that the non-canonical cleavage site of KTS1^Cmal^ is biologically active. Monobasic sites are less common in killer toxins but have been confirmed in K1 at the delta/alpha boundary and in K28 between alpha/delta and alpha/gamma. Mutagenesis data also supports the cleavage of monobasic sites in K2. A cleavage site in the middle of the gamma domain of KTS1^Cmal^ also demonstrates similar organization to K1, K2, and K28 and divides gamma into 39 and 71 amino acid peptides. All Kex2 cleavage sites of KTS1^Cmal^ are solvent exposed with 2/3 on flexible loops in relatively the same positions as K74 (Table SX).

As in the molecular models of K1, K2, K45 family killer toxins, the hydrophobic 2α helix of KTS1^Cmal^ is wrapped by a discontinuous antiparallel beta sheet composed of strands from all three domains. The alpha domain contributes beta strands 1-2β, gamma 4-7β, and beta 8-9β. Helices 1-4α interrupt the continuity of the beta sheet and separate 1-2β from 3-8β. The central hydrophobic helix 2α is also clamped by the beta domain helices 6α and 7α. As a result of these interactions, 87.3% (1721.2 Å^2^) of the surface area of 2α is buried (Table S6). The gamma domain interacts predominantly with the alpha domain, specifically the 2α helix (Figure 6). The C-terminus of helix 1α is connected to 2α by a loop that is predicted to be stabilized by the disulfide bond C69-C82, which is a common configuration in the K1, K2, and K45 families of killer toxins. Overall, the cysteine residues are all predicted to form intradomain disulfide bonds in the alpha (C69-C82) and beta domains (C326-C334). The predicted disulfide bonds in KTS1^Cmal^ and other homologs is generally inconsistent with the disulfide-linked heterodimer that was predicted for K74 [114]. There is only one K74 homolog, from the filamentous fungi *Glarea lozoyensis*, with a predicted alpha-beta interdomain crosslink but this is due to a unique cysteine configuration. Together these data suggest that the alpha/beta heterodimer of K74 and its homologs are likely held together by non-covalent interactions, as is predicted for K2, K21, KHS, and K45.

### 2.7 The K1, K2, K45, and K74 killer toxin families: Mechanistic insights

Molecular modeling of the K1, K2, K45, and K74 killer toxin families indicated a shared structural organization. Primary and secondary structure analysis has previously revealed homology between several *Saccharomyces* killer toxins, suggesting conserved tertiary structures between K1L/K1 and K2/KHS [23,41]. Other similarities between K1, K2, and K21 toxins have been based on commonalities in cellular proteins required for killer toxin resistance and their apparent mechanism of action, i.e., cell permeabilization and ion efflux. A four-domain organization is a common feature among these killer toxins, as indicated by the conserved positioning of monobasic and dibasic motifs recognized by Kex proteases. However, a shared structural layout does not directly determine a shared antifungal mechanism, as toxins such as K28 and K1 have the same domain naming convention (delta, alpha, gamma, and beta) and domain order but kill cells by fundamentally different mechanisms.

Despite low sequence identity among K1, K2, K45, and K74 families of killer toxins (7.7–32.0%), they exhibit conserved secondary and tertiary structure. Alignment of the tertiary structure models of K1, K2, K45, and K74 families of killer toxins revealed a well-defined core motif of two helices and a single sheet composed of four to seven antiparallel beta strands (Figure 7A and S6). The first helix in this motif is the central helix of the alpha domain that is buried in all tertiary structure models. The second helix from the beta domain interacts with this central helix, with the only exception being K2. The majority of beta strands are formed from sequences between the two alpha helices, with the exception of two C-terminal beta strands. A conserved structural motif is observed at the C-terminus of the beta domain, consisting of a beta strand followed by an alpha helix and then two additional beta strands. Importantly, the final beta strand is inserted between the two other beta strands to form an antiparallel beta sheet (Figure 7A and S6). Overall, this conserved core motif has an average RMSD of 4.5 Å across the K1, K2, K45, and K74 families of killer toxins, compared to 8.7 Å when comparing the entire ppTox structures.

A central helix of the K1 superfamily of killer toxins is always located within the alpha domain, which is responsible for cytotoxicity [51,106]. PSIPRED analysis predicted that the central helix in all K1 superfamily toxins interacts with membranes (Figure S6). Specifically, four of these helices are predicted to be transmembrane helices (K2, K21, KHS, and K45) and three are amphipathic pore-lining helices (K1, K1L, and K74). The differences in PSIPRED predictions could indicate mechanistic differences of membrane attack, especially as K2, K21, KHS, and K45 have additional helices and beta sheets in the alpha domain that are also predicted to interact with membranes. In the ppTox, the central helix is sequestered and buried, likely to prevent unwanted interactions with membranes and associated toxicities while enabling the correct folding and association of the alpha and beta domains. The alpha domains all share a conserved disulfide bond that pins the N-terminal end of the central helix to the C-terminal end of the preceding secondary structure element in alpha. This bond may play a conserved role in positioning or constraining the conformational flexibility of the central helix, which is important for membrane attack and pore formation. Overall, the structural conservation and biochemical characteristics of the central alpha helices support the proposed antifungal mechanism of these toxins, namely, membrane disruption likely through pore formation. The close structural similarities of these toxins also support the proposal that their mechanism of intoxication is conserved and that they represent a broader “K1 superfamily”, a naming convention based on the early discovery of K1. As such, the mechanistic insights gained from the study of K1 and K2 can be broadly applied to the wider superfamily, including hundreds of homologous sequences that remain uncharacterized.

To identify structural homologs of the killer toxins of the K1 superfamily, predicted pTox structures were used to query the protein database using DALI software. The top hit for four of the seven K1 superfamily killer toxins (K1, K1L, KHS, K45) identified structural matches with four different bacterial pilins or pseudopilins (Figure 7B). Analysis of the structural homologs that overlapped between the different killer toxins identified ten proteins that were structurally similar to four or more killer toxins. Of these ten, seven were pilins or pseudopilins from five different species of bacteria. The pseudopilin EpsI from the type 2 secretion system of the bacteria *Vibrio vulnificus* was identified as having structural homology to six of the seven killer toxins, with an average z-score of 3.0 and RMSD of 6.4 Å (Figure 7B; Table S8). The structural similarity with killer toxins is due to the characteristic pilin motif, which is an alpha helix that is partially wrapped by a four-stranded antiparallel beta sheet [115] (Figure 7). The identification of these structural homologs was unexpected, as pilins and pseudopilins are not toxins, but subunits of helical filaments that extend from bacterial cells to aid in processes such as effector secretion, DNA uptake, adhesion, and motility.

Filament formation similar to pilins is a mechanism of membrane attack and has been observed for the insecticidal cytolysins (Cyt) from the biocontrol bacterium *Bacillus thuringiensis* [116,117]. Cyt toxins have similar tertiary structures to pilins and K1 superfamily toxins, with an alpha/beta/alpha sandwich organization [118,119]. K1 is also capable of oligomerizing into large complexes or aggregates, but has been considered only as evidence of membrane pore formation [48,85,86]. For Cyt toxins, there are two models for their cytotoxic mechanism, one that is predicted to resemble the action of a detergent and another that involves the formation of membrane pores [120–122]. Importantly, filament formation and pore formation may not be mutually exclusive mechanisms of membrane attack and could depend on the biological properties of the target membrane and toxin concentration [120,123]. Whether the structural similarity of K1-superfamily proteins to pilins is biologically relevant or just a coincidence of their common structural organization remains to be further investigated.

In addition to pilins, DALI also identified two notable homologs to the K1 superfamily: the antifungal protein ginkbilobin-2 (Gnk-2) from the *Ginkgo biloba* tree and the salt-mediated killer toxin (SMKT) from *Millerozyma farinose* [124,125]. Gnk-2 was more structurally similar to K1L (z-score 3.5), while SMKT was more similar to KHS, but with a low z-score of 2.1. Notably, homology between K2 and SMKT has been previously reported, with shared secondary and tertiary structures and functional motifs (Figure 7B) [107]. The RMSD of SMKT to the pTox model of K2 was 6.2 Å, but it was a closer structural match to the K1 superfamily toxins, specifically K1 (5.2 Å), K45 (5.2 Å), and KHS (5.9 Å). Both Gnk-2 and SMKT are alpha/beta sandwich proteins belonging to a diverse family of cytotoxic proteins and lectins (Pfam: PF01657, PF21414, PF21415). Like K1, the antifungal mechanism of SMKT is thought to involve membrane attack and to permeabilize artificial liposomes, consistent with pore formation. Unlike K1, SMKT does not require cell wall or membrane receptors to cause intoxication, and can directly interact with membranes of various compositions. SMKT has a domain organization of alpha/gamma/beta, similar to the K1 superfamily, as well as an amphipathic alpha helix in the alpha domain. The SMKT alpha domain stably associates with fungal membranes, whereas the beta domain is only loosely associated. This function mirrors that of the proposed mechanism of K1, where the hydrophobic alpha domain alone is toxic to yeasts and predicted to associate with membranes. Similar to SMKT, the K2, K45, and K74 toxins are predicted to lack interdomain disulfide linkages, and association of the alpha and beta domains is stabilized by non-covalent interactions. Despite variations in sequence and size, their conserved structural features and domain organization suggest a common mechanism of action across killer toxins, most likely centered on membrane disruption. However, additional structural analysis presented later in this manuscript suggest that SMKT may also be closely related to the Klus family of killer toxins.

### 2.8 The Klus killer toxin family

Klus was discovered during a screen for *S. cerevisiae* killer yeasts associated with 110 spontaneous fermentations of grapes in the Ribera del Guadiana region of Spain [42]. The screen isolated 423 strains of killer yeast, with the majority being of type K2. However, 7% of the killer yeasts exhibited a unique spectrum of antifungal activity that was due to the production of the killer toxin Klus [42]. Klus killer yeast strains were also able to inhibit the growth of K1, K2, and K28 killer yeasts and were also resistant to their own toxins. As with other *Saccharomyces* killer toxins, Klus is encoded by a dsRNA satellite named Mlus, which has been identified in *S. cerevisiae* from other regions of Spain and around the world [94]. As expected, curing of Mlus from *S. cerevisiae* resulted in the loss of killer toxin production. Klus is most active at pH 4.0 to 4.7 and between 28°C and 30°C. Klus was also previously shown to share sequence homology to an extracellular protein of unknown function encoded by the gene YFR020W in *S. cerevisiae* [126].

KHR is a genomically encoded toxin in *S. cerevisiae*, isolated from Japanese wineries in Yamanashi Prefecture, and has been identified in many strains of *S. cerevisiae* [102]. KHR is thermostable as it retains its antifungal activity from 0°C to 40°C and between pH 5.0 to 6.0 [38]. Mature extracellular KHR has an observed molecular weight of ∼20 kDa, indicating that it is likely post-translationally modified by proteolytic cleavage and is close to the theoretical molecular weight of an alpha and beta domain heterodimer (18.4 kDa). It has antifungal activity against *N. glabratus*, *S. cerevisiae*, and *Kluyveromyces lactis*.

#### 2.8.1 The Klus killer toxin family: Domain organization and posttranslational modification

Klus has three predicted dibasic Kex2 cleavage sites, resulting in a classic four-domain configuration (24-67, 68-98, 99-167, and 168-242), similar to other known killer toxins [42]. This organization includes a predicted signal sequence cleavage site after amino acid 23, which would allow entry into the secretory system. However, the location of the six cysteine residues in the last two domains of the protein represents a unique configuration compared to other killer toxin families, which typically have most of their cysteines in the second and last domains. Klus is also predicted to have a transmembrane region that includes a portion of the 1α helix in the third domain from the N-terminus. Based on these data and additional data from modeling and comparison with empirically determined structural models, the domain organization for Klus is predicted to be gamma, delta, alpha, and beta.

Overall, KHR has a similar pattern of secondary structure compared to Klus, but is 54 amino acids longer with additional structural elements and one additional Kex2 cleavage site. The positioning of the proteolytic cleavage sites suggests that KHR has four domains between amino acids 22-77, 78-130, 131-183, 184-296, and a signal sequence cleavage site after residue 21. As is the case for Klus, there is also a predicted transmembrane region in the third domain from the N-terminus of KHR. KHR also has 11 cysteine residues, which, like Klus, are mostly positioned in the third and fourth domains, supporting their assignment as the alpha and beta domains in the C-terminal half of the toxin. Although KHR was identified as a sequence homolog of Klus, and the position of the alpha and beta domains is the same as Klus, the organization of the secondary structure suggests that the gamma and delta domains are reversed in KHR relative to Klus (delta/gamma/alpha/beta). Further justification for the unique rearrangement of the structural domains of KHR and Klus is presented below.

#### 2.8.2 The Klus killer toxin family: Molecular modeling

The tertiary structure model of Klus is a globular protein composed of a central 1α helix (alpha domain) positioned next to a shorter 2α helix (beta domain) that is offset from parallel by ∼45 degrees. The 1α helix is amphipathic and predicted by PSIPRED to be pore-lining, consistent with an ionophoric mechanism of action. A five-stranded discontinuous beta sheet composed of beta strands from the gamma, alpha, and beta domains with the 2α helix creates a pocket that buries 1386.0 Å² (59.3%) of 1α (Table S6). The beta sheet consists of two pairs of discontinuous beta strands (3-4β and 5-6β) that are aligned in an antiparallel configuration. Beta strand 1β is aligned in parallel on the outer edge of the beta sheet, adjacent to strand 5β. The inclusion of 1β into the beta sheet that wraps the hydrophobic helix justifies the assignment of the N-terminal domain as gamma. The cradling of the pair of helices by the beta sheet positions 1α to interact with 3-4β and 2α with 1β/5-6β. Moreover, 2α is clamped between the 2β antiparallel sheet and 1α, burying the helix more extensively than 1α. The opposite side of the beta sheets, relative to 1α and 2α, is a tangle of unstructured protein sequence, consisting of the N-terminus, C-terminus, and a 36 amino acid loop that straddles the delta and alpha domain boundary. Predicted disulfide bonds in Klus connect the N-terminal end of 1α and the C-terminal end of 2α (C114-C188) and the C-terminal end of 1α to the N-terminal end of 2α (C141-C162). A third pair of disulfide bonds stabilizes an unstructured loop at the C-terminal end of Klus, a similar linkage was observed for the majority of killer toxins in the K1 family. Proteolytic processing of the pTox by Kex2 would mean that C114-C188 would tether the alpha and beta domains, whereas C141-C162 would pin the 3-4β sheet to 1α. This organization of disulfide bonds supports the unique domain organization of Klus, and the mature toxin being a disulfide bond linked heterodimer after the cleavage and dissociation of the N-terminal delta and gamma domains.

KHR is also a globular protein with a similar overall organization to Klus, composed of two helices: 3α (alpha domain) and 4α (beta domain), which are positioned ∼45 degrees from parallel. The 3α helix of the AlphaFold2 model is a 17-residue helix, but unravels to a predominantly unstructured coil and smaller helix after MD simulations. The remaining helical structure has a buried surface area of only 380.5 Å^2^ (Table S6). KHR has a domain ordering of delta, gamma, alpha, and beta, with the latter three domains contributing to the organization a six-stranded discontinuous beta sheet formed from three pairs of antiparallel beta sheets (3-4β, gamma; 7-8β, and 9-10β, beta). The inclusion of 3-4β into the beta sheet justifies the assignment of the gamma domain as the second domain from the N-termini. Unlike Klus, the extensive beta sheet does not include strands from the alpha domain (5-6β). The opposite side of the beta sheet to the 3α and 4α helices is a region of extensive elaboration compared to Klus, featuring small helices and sheets, as well as large unstructured loops from the delta, gamma, and alpha domains. Curiously, the alpha domain contains a dibasic cleavage site that would split the domain in two but is bridged by a predicted disulfide bond between C138 and C151 (Table XX disulfide). Importantly, in KHR models, there was no indication of interdomain disulfide bonds as was predicted for Klus and other more well-studied killer toxins.

To better compare Klus and KHR to existing empirical protein structures, models of the mature toxin structures consisting of alpha and beta domains were constructed using AlphaFold2 multimer prediction mode. These heterodimeric structures had high confidence (average pLDDT 71.0 (KHR) and 72.4 (Klus)) and were stable over a 1 µs MD simulation. The high confidence of the predicted mature toxins contrasts the low confidence of other *Saccharomyces* killer toxins that had an average pLDDT score of 45.4 (Table table XX<u>?</u>). During MD, Klus and KHR structures quickly achieved similar conformations within 10 ns that stabilized around 0.3 nm RMSD (Figure S4). The mature Klus and KHR did not recapitulate the exact structural organization of their respective pTox as they folded new regions of secondary structure and rearranged existing structural elements. These included the extension of the previously small 3α helical region in KHR to a long 54 amino acid helix (compare Figure 8G to 9). In Klus, 1β of the gamma domain was replaced with beta strands folded from an unstructured loop of the alpha domain. For KHR, beta strands 3-4β of the gamma domain were replaced by 5-6β from the beta domain. Disulfide bonds in mature Klus were unchanged from pTox and indicated that the mature heterodimer is bonded between the alpha and beta domains by C114-C188. For mature KHR, the configuration of disulfides was significantly different from pTox in that only C248-279 was maintained when comparing the predicted pTox and mature toxins. Alternative disulfide pairings were predicted that created two interdomain bonds between alpha and beta domains (C138-C294 and C143-C230) and one intradomain bonds in the beta domain (C189-C219).

**Figure 8.**
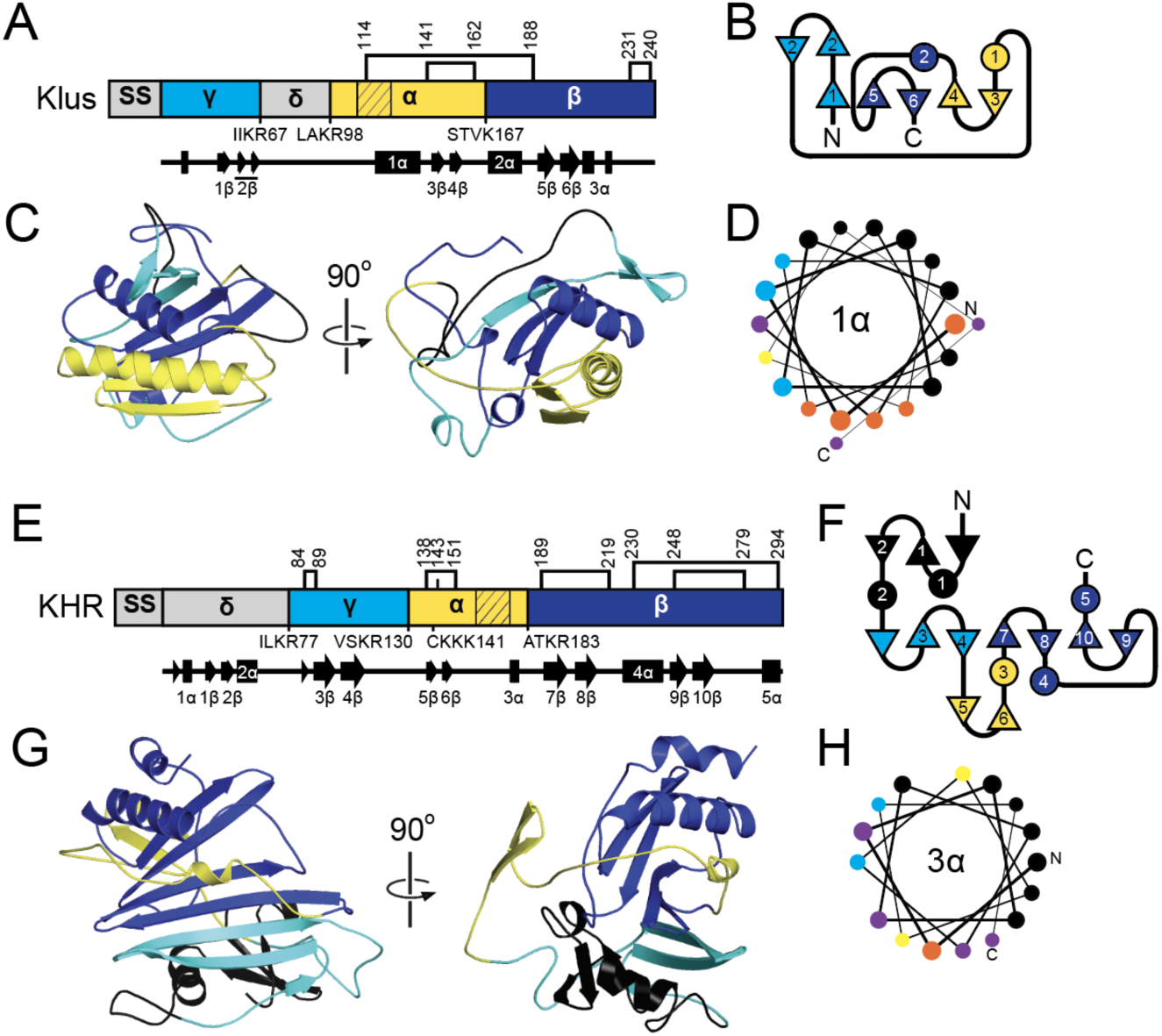
Secondary and tertiary structure models of Klus family killer toxins. (A, E) Domain diagrams of Klus and KHS ppTox indicating the sites of proteolytic processing that define the delta, alpha, gamma, and beta domains. The positioning of Kex protease cleavage sites is indicated below the domain diagram, with the four amino acids preceding the cleavage point illustrated. Cysteine residues are indicated by lines and numbers above the domain diagram, with connections between them representing predicted disulfide bonds. Hatching in the diagram represents amino acid sequences predicted to form transmembrane or pore-forming structures. Secondary structure, with numbered alpha helices and beta strands, is represented by arrows and rectangles, respectively, below the domain diagram. (B, F) Two-dimensional schematic of the relative secondary structure organization of Klus and KHR, with helices represented as circles and beta strands as triangles. N, amino-terminus domain, C, carboxyl-terminus. (C, G) Tertiary structure model of killer toxin pTox colored by domains as depicted in panels A. (D, H). Helical wheel diagram of 2α helix, with hydrophobic (black), positively charged (yellow), negatively charged (blue), polar (purple), and aromatic (orange) amino acids. The line thickness between amino acids (from thick to thin) represents the progression of the sequence from the N-terminus to the C-terminus. During the MD simulation, the 3α helix unfolded; therefore, panel D represents the relaxed AlphaFold2 predicted helix before it was subjected to molecular dynamics.

**Figure 9.**
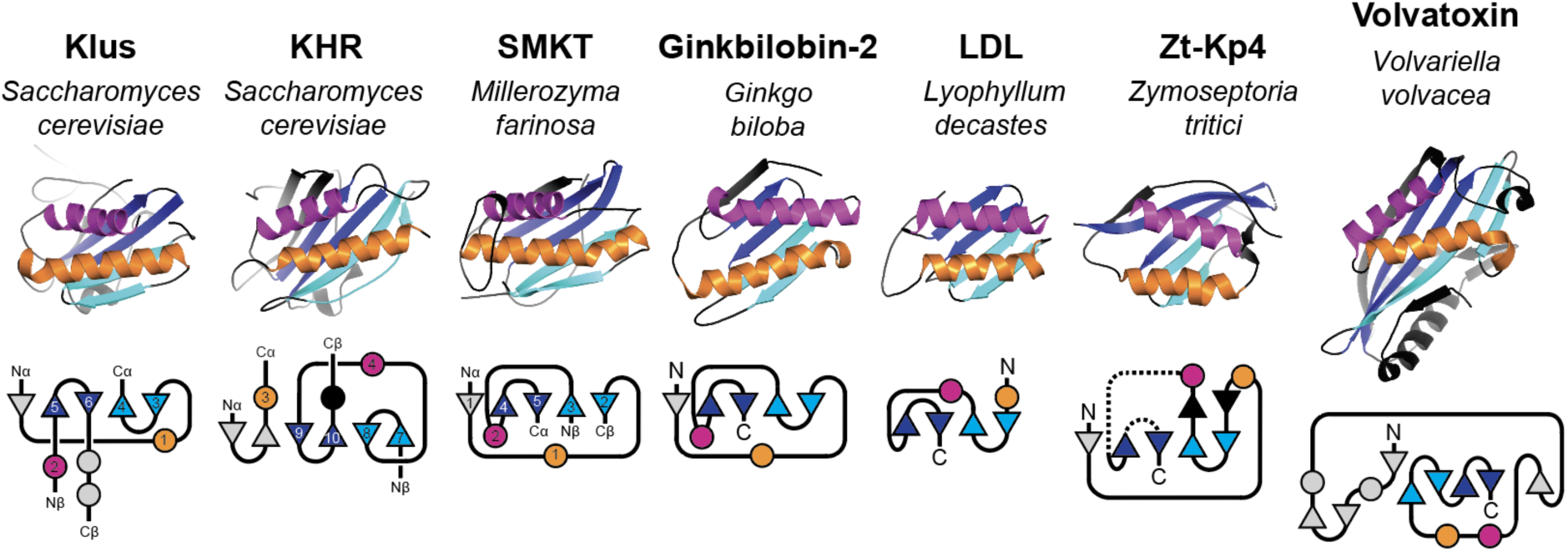
Structural homologs of the Klus family of killer toxins. Empirically determined tertiary structures of fungal proteins with homology to Klus and KHR. Secondary structure elements of mature Klus and KHR are labeled based on pTox (Figure 8) and unlabeled secondary structures in the models are assembled from disordered regions of the pTox. Secondary structure is colored to emphasize beta strand pairs and helices common to all proteins. Below the tertiary structures are two-dimensional schematic of the relative secondary structure organization of Klus and KHR and their structural homologs with an emphasis on the relative organization of the beta sheet. Helices are represented as circles and beta strands as triangles, N - amino-terminus domain, C - carboxyl-terminus, Nα/Cα - alpha domain amino/carboxyl-terminus, Nβ/Cβ - beta domain amino/carboxyl- terminus.

#### 2.8.3 The Klus killer toxin family: Mechanistic insights

Comparison of the tertiary structures of Klus and KHR revealed average RMSD values of 5.7 Å (pTox) and 4.1 Å (mature), confirming the observed sequence similarities. Similar to the K1-superfamily, querying the DALI server with the mature and ppTox models identified homology to alpha/beta sandwich protein families (Pfam: PF01657, PF21414, PF24145, and PF09044). In particular, the PF01657 family includes a large (30,000+) subfamily of plant-specific proteins that contain the Domain of Unknown Function 26 (DUF26), including cysteine-rich receptor kinases and secreted cysteine-rich proteins. These proteins are used for responding to environmental and biotic stressors, including bacteria and fungi [127]. Despite little to no sequence identity to Klus and KHR, many of the structural homologs had a pair of helices bound to one side of a four-to-six stranded antiparallel beta sheet (Figure 9). Consistent with the cytotoxic activities of Klus and KHR, many structural homologs were identified that are cytotoxic to fungi, including the antifungal proteins Gnk-2, SMKT, and Zt-KP4 [46,124,125]; other homologs are toxic to cells of higher organisms (Lyophyllum decastes lectin (LDL), and Volvatoxin) [128,129] (Figure 9). Compared to prior pTox models of K2, K1L, and KHS that were previously judged to have some structural similarity to SMKT (RMSD = 5.2-6.2 Å), the structural alignment of mature Klus and KHR to the empirical SMKT structure resulted in lower RMSD values (3.2 Å and 4.1 Å, respectively).

Despite the close tertiary structure match of SMKT to Klus and KHR, the domain order of SMKT is more similar to other canonical killer toxins (delta/alpha/gamma/beta) (Figure 10A) [130]. However, it was unclear if the pTox structure of SMKT would match better with the K1 superfamily or Klus family toxins. To compare the structures of these immature toxins, the pTox structure of SMKT was determined by AlphaFold. The predicted configuration of the SMKT pTox overlapped almost perfectly with the alpha/beta heterodimer of the mature SMKT (RMSD 0.28 Å), but with the replacement of pTox β5 from the gamma domain with a beta strand folded from an unstructured loop in the alpha domain of the mature SMKT [107] (Figure 10B). The close similarity in the tertiary structures of SMKT and its pTox meant that Klus and Klus also had a similar tertiary structure to SMKT pTox (RMSD, 3.6 Å and 4.2 Å) (Table S9). SMKT pTox also had structural homology to the K1-superfamily but with a higher RMSD compared to Klus and KHR (5.2-5.3 Å; Table XX). As was observed in the pTox models of the K1 superfamily, Klus, and KHR, the SMKT pTox extensively wrapped a central helix (1α) with a large eight-stranded antiparallel beta sheet assembled from beta strands from the alpha (1β), beta (6-8β) and gamma (2-5β) domains (Figure 10B and C). The similar structures reflect the importance of the respective gamma domains in pTox folding and the burying of the centrally positioned transmembrane helix in SMKT and *Saccharomyces* killer toxins (Figure 10D). Killer toxin immunity is another known function of the killer toxin gamma domain of K1, but differences in the Klus and KHR domain order and the positioning of gamma domain beta strands could indicate a non-canonical immunity mechanism. Moreover, it is unclear whether SMKT encodes functional immunity, as ectopic expression by *S. cerevisiae* results in a suicidal phenotype [131]. Overall, the tertiary structures of Klus and KHR are most similar to SMKT, despite their unusual domain organization, but it remains to be determined whether the structural homology can be used to infer functional similarities.

**Figure 10.**
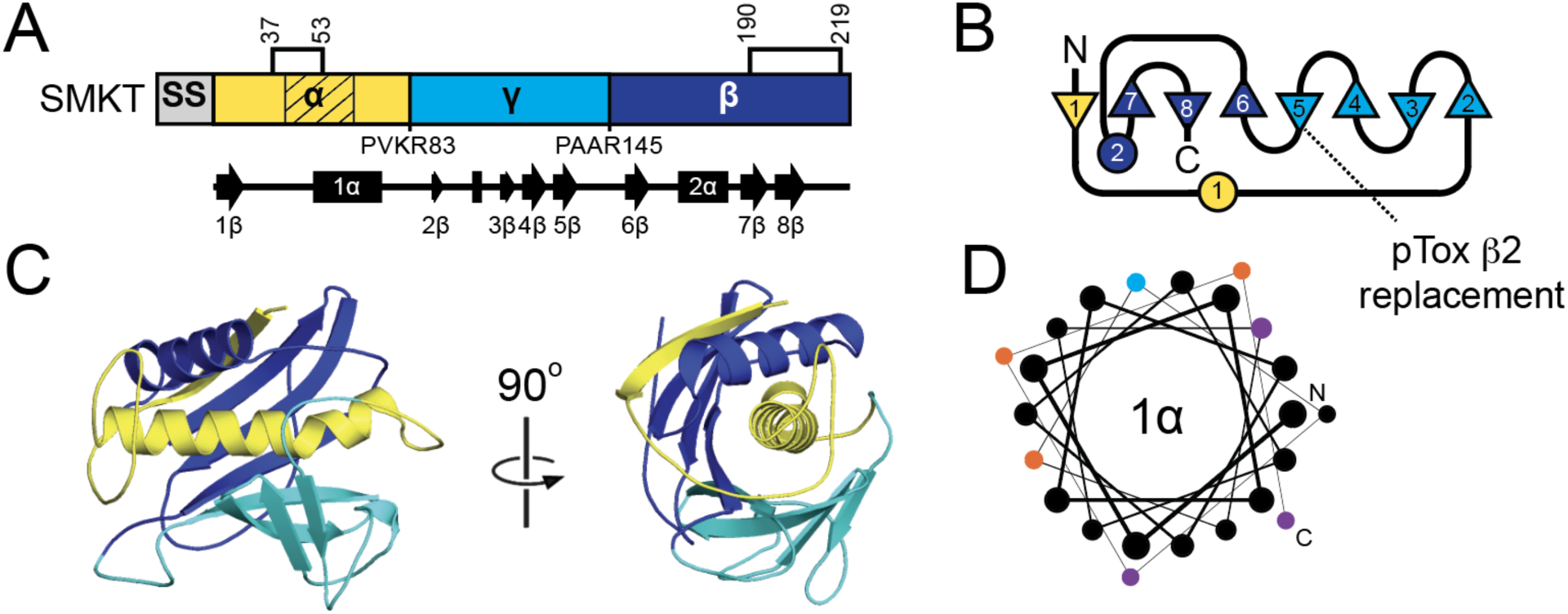
Molecular modeling of the SMKT pTox. (A) Domain diagrams of SMKT ppTox indicating the sites of proteolytic processing that define the alpha, gamma, and beta domains. The positioning of Kex protease cleavage sites is indicated below the domain diagram, with the four amino acids preceding the cleavage point illustrated. Cysteine residues are indicated by lines and numbers above the domain diagram, with connections between them representing predicted disulfide bonds. Hatching in the diagram represents amino acid sequences predicted to form transmembrane or pore-forming structures. Secondary structure, with numbered alpha helices and beta strands, is represented by arrows and rectangles, respectively, below the domain diagram. (C) Two-dimensional schematic of the relative secondary structure organization of Klus and KHR, with helices represented as circles and beta strands as triangles. N, amino-terminus domain, C, carboxyl-terminus. (D) Tertiary structure model of SMKT pTox colored by domains as depicted in panel B. (E) Helical wheel diagram of 2α helix, with hydrophobic (black), negatively charged (blue), polar (purple), and aromatic (orange) amino acids. The line thickness between amino acids (from thick to thin) represents the progression of the sequence from the N-terminus to the C-terminus.

**Figure 11.**
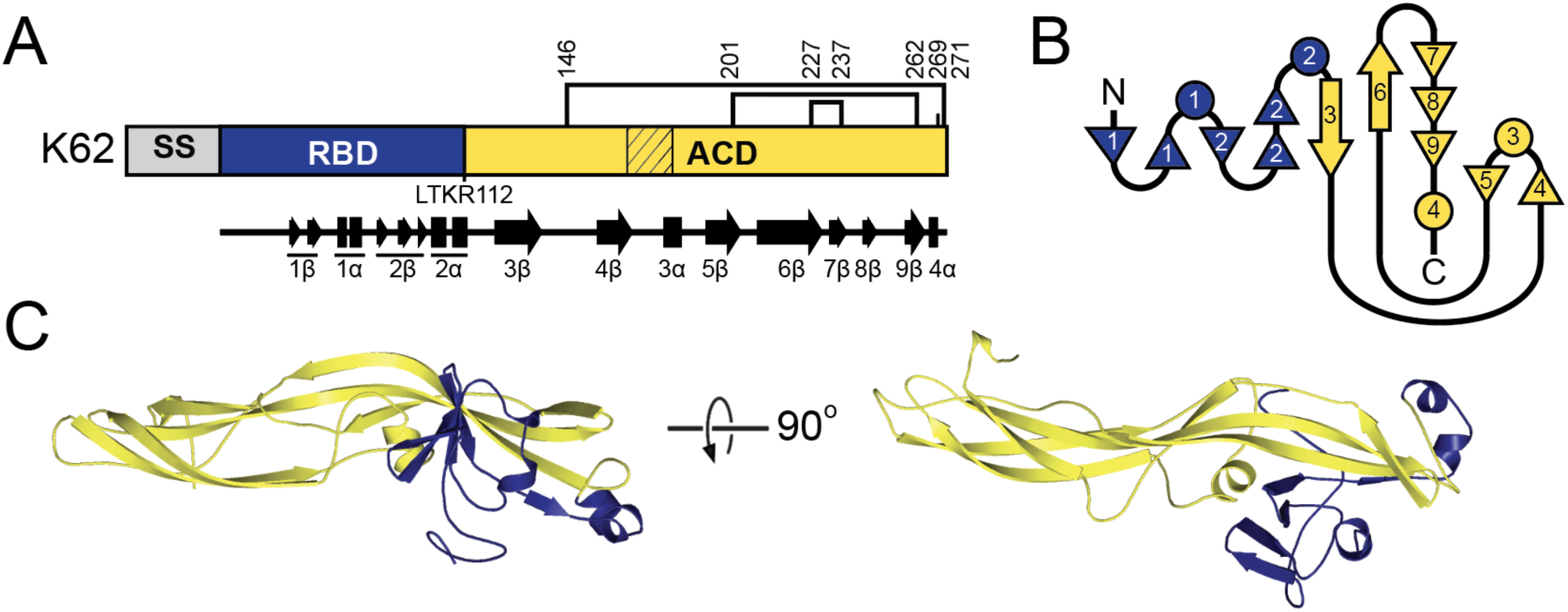
K62 Structural Model. (A) Domain diagram of ppTox K62 representing the N-terminal signal sequence (SS), putative receptor bonding domain (RBD) and aerolysin core domain (ACD). The domain boundaries are defined by a predicted signal sequence cleavage site, and a single site of proteolytic processing based on the position of a dibasic (KR) motif. Disulfide bonds are represented as lines between cysteine residues. The hatched region indicates a predicted transmembrane sequence. Below the domain diagram is a representation of pTox K62 secondary structure with alpha helices and beta sheets shown as rectangles and arrows, respectively. (B) Two-dimensional schematic of the relative secondary structure organization of K28 with helices represented as circles and beta strands as triangles. Number of the secondary structure elements is consistent with panel A. (C) Tertiary structure model of killer toxin pTox colored by domains as depicted in panel A.

The intoxication mechanism of SMKT is thought to involve disruption of biological membranes, providing possible insights into the antifungal mechanisms of Klus and KHR [132]. SMKT also has structural homology to VVA2 and Cyt toxins, which attack membranes. Membrane interaction by these toxins appears to be independent of cell surface receptors and interacts directly with membrane lipids. Cyt toxins cause specific ion leakage, leading to a model of pore formation as a possible cytotoxic mechanism. Amphipathic helices in the alpha domains of SMKT and Klus family proteins are positioned in the same location as amphipathic helices in Cyt and VVA2 (Figure 9). In VVA2, this helix is important for oligomerization and toxicity, but not for membrane interaction [133]. Indeed, C-terminal beta strands of VVA2 and Cyt toxins stably associate with membranes, suggesting the formation of a beta-barrel pore. The membrane association of the C-terminal domains of Cyt and VVA2 contrasts with SMKT, which stably associates its N-terminal alpha domain with the membrane fraction of intoxicated yeasts [132]. A model of alpha domain membrane attack would be more similar to the K1 superfamily of toxins. Additionally, the C-terminal beta domain of SMKT appears to only be loosely associated with membranes, which would contradict a model of C-terminal domain beta barrel formation [132]. In contrast to pore formation, VVA2 and Cyt toxins are also capable of oligomerization into filaments (as discussed in the K1 superfamily section). Cyt toxin filaments are thought to cause membrane delamination and the collapse of large liposomes to smaller vesicles and lipid aggregates. A similar dramatic destruction of liposomes has also been observed by SMKT, but it is unclear if it requires toxin oligomerization for this mode of action.

Other mechanistic insights into the antifungal activities of Klus and KHR can also be drawn from the KP4-like killer toxin from the wheat pathogen *Zymoseptoria tritici* (Zt-KP4-1) [46]. This antifungal toxin is a close structural homology to the well-studied antifungal protein KP4 from the corn-smut fungus *Mycosarcoma maydis* (syn. *Ustilago maydis*) and toxin homologs from other fungal pathogens [45,134]. However, unlike SMKT, KP4 is not predicted to attack fungal membranes but instead blocks L-type voltage-gated calcium channels in both fungal and mammalian cells [135,136]. KP4 and its homologs also inhibit plant root growth [45,137]. Consistent with the interference of ion homeostasis, high concentrations of calcium can protect cells and plants from intoxication by KP4 and its homologs. However, the precise molecular interactions by which KP4 targets and disrupts calcium channels remain unresolved, in addition to whether KP4 and its homologs interact directly or indirectly with calcium channels.

Other alpha/beta sandwich proteins with structural similarity to Klus and KHR include the α-galactosyl binding lectin from the mushroom *Lyophyllum decastes* (LDL), the glycan binding protein Y3 from *Coprinus comatus*, and Gnk-2. These proteins are toxins with activities against higher eukaryotic cells (LDL and Y3) and fungal cells (Gnk-2) [138–140]. How these toxins cause damage to cells is unknown, but all bind specific carbohydrates and have defined surface-binding pockets on the same face of the tertiary structure close to the C-terminal end of the second helix [139–141]. Carbohydrate binding suggests the possibility of similar motifs in Klus and KHR based on their conserved tertiary structures. The potential for carbohydrate binding by Klus and KHR would be similar to other killer toxins that have carbohydrate cell wall receptors, i.e. glucans and mannan. However, SMKT is also a structural homolog of Klus and KHR and appears not to require cell wall binding to cause cytotoxicity.

The structural similarity of Klus and KHR to other alpha/beta sandwich proteins offers insight into their potential mechanisms of action. Members of this protein family are known to kill cells by disrupting membranes or interfering with ion homeostasis, indicating that the alpha/beta sandwich fold can support diverse cytotoxic functions. However, the exact mechanisms underlying these activities remain unresolved, and it is unclear whether observed functional differences reflect true mechanistic divergence or variation in experimental systems. A key goal for future research is to determine whether these proteins share a common mode of intoxication or if substantial functional diversification has occurred around a conserved tertiary structure.

### 2.9 The K28 killer toxin family

The first K28 killer yeast was isolated from grapes and identified by screening 163 yeasts collected by the Johannes Gutenberg-Universität Mainz for the killer phenotype [49]. Initially described as being produced by *S. cerevisiae,* but was later confirmed to be *S. paradoxus*. The antifungal activities of K28 appeared to be different from those of other killer toxins, and K28 killer yeasts were uniquely sensitive to K1 and K2, suggesting a distinct toxin type [50]. K28 is encoded on a satellite dsRNA named M28, and curing of the dsRNA by cycloheximide caused the concomitant loss of the K28 killer phenotype [142]. Determination of the M28 genetic sequence identified the K28 gene but revealed no sequence homology to any other protein in the database [143]. The antifungal activity of K28 is optimal at pH 5.8, which is less acidic than optimal conditions for other killer toxins [49,50]. K28 is also more tolerant of higher temperatures, remaining active at 40°C for up to one hour.

#### 2.9.1 The K28 killer toxin family: Domain organization and posttranslational modification

Like all other *Saccharomyces* killer toxins, K28 is trafficked through the secretory system to enable maturation and extracellular localization. After translation, exportation of the ppTox to the ER is directed by an N-terminal signal sequence that is cleaved after G36. Mutagenesis of G36 and N-terminal sequencing of the 42 kDa K28 pTox confirmed the location of signal sequence cleavage [143,144]. Purification of the extracellular K28 resulted in the detection of a ∼16 kDa disulfide-linked heterodimer consisting of two polypeptides that were named alpha (10.5 kDa) and beta (11.0 kDa). N-terminal sequence analysis of extracellular K28 confirmed that alpha and beta domains begin at amino acids 50 and 246, respectively [143]. The identification of alpha and beta led to the definition of the four functional domains of K28 pTox (delta, 36-49; alpha, 50-149; gamma, 150-245; beta, 246-344), based on the presence of dibasic motifs that are targeted by the Golgi-resident endopeptidases Kex1 and Kex2 [144]. Consistent with the major role of Kex2 in the proteolytic processing of K28, a kex2Δ null strain does not produce active K28 and is unable to cleave pTox to release the gamma and delta domains from pTox. Mutation of dibasic sites at the predicted domain boundaries prevented the expression of active K28 and resulted in incomplete pTox cleavage. Although not initially thought to be essential for K28 activity, Kex1 removes a critical C-terminal arginine residue to reveal an ER retention motif “HDEL” that is essential for efficient K28 retrograde trafficking into K28 susceptible cells [145,146].

With only a single cysteine residue present in the alpha domain (C56), it was deemed essential for heterodimer formation by mutagenesis. Of the four additional cysteines present in beta, only C340 was essential for K28 toxicity, as mutagenesis of the remaining cysteines to tryptophan (C292W, C307W, C333W) had no apparent effect on K28 activity. However, a more extensive follow-up study identified that C56-C333 was the more likely interdomain disulfide bond, with the remaining beta domain cysteine residues important for the toxicity and stability of K28 [147].

#### 2.9.2 The K28 killer toxin family: Antifungal mechanism and immunity

The antifungal mechanism of K28 causes cell cycle arrest at G1/S, arresting DNA synthesis through an unknown mechanism [83,148,149]. Cell targeting relies on the interaction of the beta domain with a primary cell wall receptor and secondary cell membrane receptor [54,145,146]. K28 interacts with yeast cell wall mannans, specifically the short, branching mannose chains attached to the 1,6-linked mannose main chain, which are critical for K28 interaction with the cell wall [150,151]. Moreover, mutants lacking functional mannosyl transferases (i.e., *MNN1*, *MNN2*, and *MNN5*), which attach mannose residues to the main chain via 1,2 and 1,3 glycosidic bonds, are resistant to K28, but not to killer toxins K1 and K2, which use cell wall 1,6-beta-D-glucan as their primary cell wall receptor [15,152,153].

Following the binding of the cell wall mannans, K28 is translocated to the cell membrane, where the beta domain interacts with a secondary cell surface receptor, Erd2, via a C-terminal HDEL motif [145]. This interaction allows the endocytosis and retrograde transport of K28 to the ER. K28 was shown to require the protein disulfide isomerase (Pdi1) to prevent toxin inactivation by unproductive oligomerization in the more neutral pH of the ER [147] . K28 then crosses into the cytosol from the ER, likely using the Sec61 translocase complex, via a mechanism distinct from the ER-associated protein degradation (ERAD) pathway [154]. In the cytoplasm, the disulfide bond linking the alpha-beta heterodimer is reduced, and the beta domain is ubiquitinated and degraded. The alpha domain moves to the nucleus, where it induces cytotoxicity by arresting cells at G1/S phase through an unknown mechanism [83,148,149].

The mechanism of K28 immunity depends on the K28 ppTox, which is analogous to other killer toxins but has a unique mechanism. Similar to K1, the K28 alpha domain is required for immunity and requires a C-terminal extension to fully protect from exogenous K28 [155]. For example, expression of the alpha domain confers partial immunity to exogenous killer toxin, whereas expression of alpha and gamma confers full immunity; the exact sequence of gamma is not essential for immunity [155]. K28 immunity is based on a fraction of ppTox being present in the cytosol to intercept mature K28 during its transit from the ER to the nucleus (via the cytosol). Complex formation between mature and ppTox K28 in the cytosol is thought to result in ubiquitination and the degradation of K28 by the proteasome.

In addition to ppTox-derived immunity, there is also a dedicated Killer Toxin Defense factor named *KTD1* that protects cells from K28 [20]. Alleles of *KTD1* in *S. cerevisiae* impart different levels of K28 resistance and elevated rates of non-synonymous substitutions in the gene, suggesting that it is under positive selection, a signature of a genetic arms race, likely with K28. The mechanism of *KTD1* protection differs from ppTox immunity, as it appears to intercept K28 in the endosomal trafficking system, preventing retrograde translocation to the nucleus. Indeed, Ktd1 requires the conserved oligomeric Golgi (COG) complex for proper antitoxin function and correct localization [19] . Signatures of positive selection in *KTD1* would suggest that there is a direct interaction with K28, and this may reroute the toxin to the vacuole, preventing it from reaching the nucleus and arresting cells in G1/S phase.

**Figure 10.**
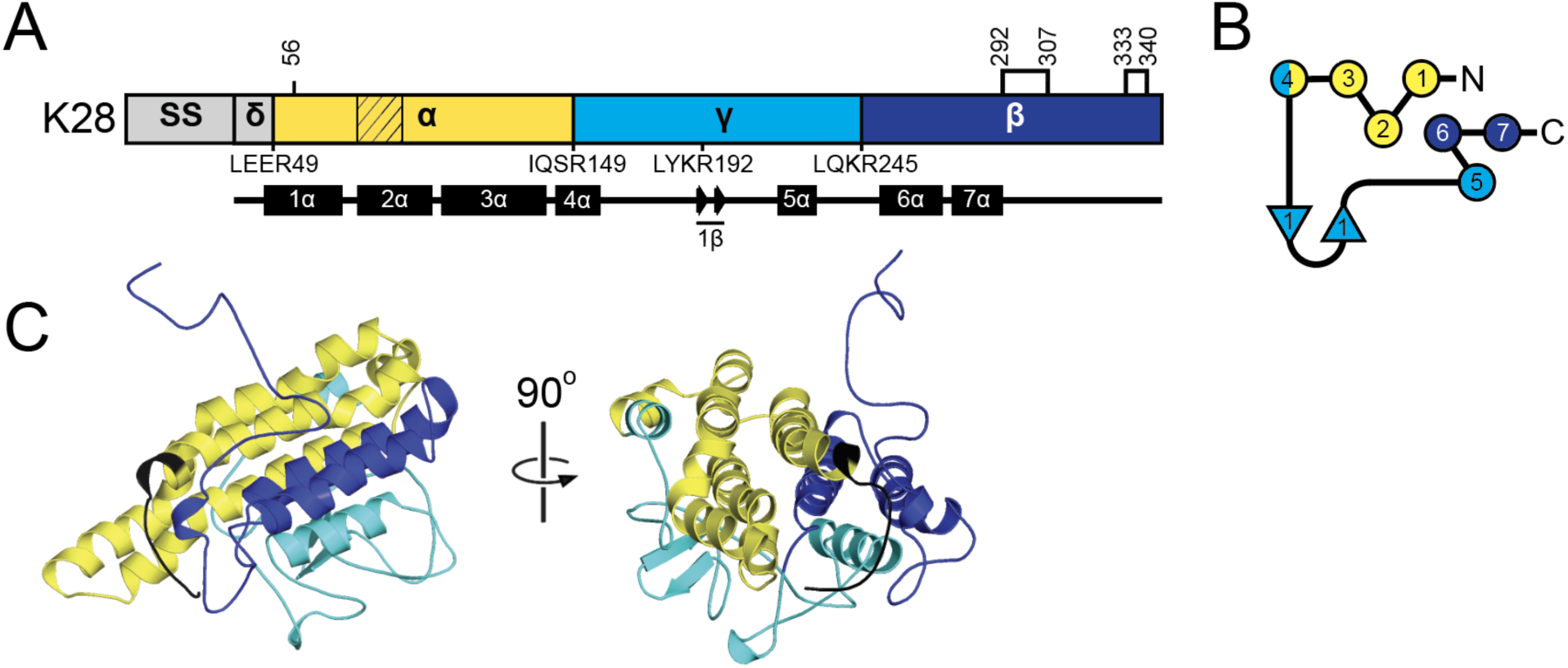
K28 Structural Model. (A) Domain diagrams of K28 ppTox indicating the sites of proteolytic processing that define the delta, alpha, gamma, and beta domains. The positioning of Kex protease cleavage sites is indicated below the domain diagram, with the four amino acids preceding the cleavage point illustrated. Cysteine residues are indicated by lines and numbers above the domain diagram, with connections between them representing disulfide bonds predicted in the AlphaFold2 model. Hatching in the diagram represents an amino acid sequence predicted to be transmembrane or pore-forming lining by MEMSTAT. Secondary structures, including numbered alpha helices and beta strands, are represented by arrows and rectangles, respectively, below the domain diagram. (B) Two-dimensional schematic of the relative secondary structure organization of K28 with helices represented as circles and beta strands as triangles. Number of the secondary structure elements is consistent with panel A. (C) Tertiary structure model of killer toxin pTox colored by domains as depicted in panel A.

#### 2.9.3 The K28 killer toxin family: Molecular modeling

The molecular model of K28 had a low pLDDT confidence with an average of 35.0, a minimum of 19.0 and a maximum of 60.4. Modelling with AlphaFold3 did not improve the confidence in the predicted structure (pTM = 0.21). The two predicted structures had a similar overall organization, but with a deviation in the alignment of the polypeptide backbone (RMSD 9.7 Å). MD simulation of the AlphaFold2 model also indicated structural instability, with observed flexibility in the gamma domain, the N-terminal and C-terminal sequences, and helix 3α (Figure S2). The lack of sequence homologs to K28 prevented additional modeling that had previously been successful in obtaining more confident structural models of the K74 family of toxins. The low confidence of the model means that the following insights into the structure and function of K28 should be treated with caution.

The K28 model is predominantly alpha helical (51.1%), assembling into a helical bundle composed of seven helices orientated along the same axis. Flexible loops of varying length enable the antiparallel packing of the helices. The alpha domain contains helices 1-4α, the gamma domain includes part of 4α and 5α, and the beta domain contains helices 6-7α. Helices 2α and 6α are in the core of the structure, surrounded by other helices, with 2α of the alpha domain being predicted to be transmembrane. Domain boundaries defined by Kex cleavage sites are located on exposed, flexible loops that would be readily accessible for proteolysis during K28 maturation. The C-terminal tail of K28 contains the HDELR motif that is essential for retrotranslocation from the cell surface to the cytoplasm. In the tertiary structure model, this motif is exposed at the end of a flexible linker, making it accessible to the action of the Kex1 carboxypeptidase, which removes the terminal arginine, allowing for recognition by the Erd2 receptor.

The K28 model does not predict a disulfide bond between the alpha and beta domains as reported in the mature toxin (C56-C333). Instead, the four cysteines in the beta domain form a pair of intradomain disulfide bonds with C56 remaining unbonded. MD simulations revealed that the K28 model is flexible and that C56 moved from 51.7 Å to 20.1 Å relative to C333 during simulations. The observed structural flexibility could allow for the linkage of the alpha and beta domains via a C56-C333 disulfide bond, as well as alternative disulfide configurations. The flexibility of the cysteine pairing in K28 has been shown to trigger unproductive oligomerization at neutral pH [147] (Table S7). Disulfide rearrangement also enables the release of the alpha domain into the cytoplasm, a critical step for intoxication by mature K28. As immature cytoplasmic K28 is essential for immunity to exogenous mature toxin, the current model may more closely represent the structure of the reduced immature ppTox. The current structural model of pTox K28 provides insights into the structural flexibility of immature K28 and an alternative conformation that might be relevant for K28 immunity, maturation, export, retrograde transport, or intoxication.

#### 2.9.4 The K28 killer toxin family: Mechanistic insights

Analysis of the K28 tertiary structure model using DALI identified 34 protein structures with z-scores greater than 4.0, which are functionally diverse and include ferritins, saccharide translocases, interleukins, and motor proteins. The top hit from DALI analysis, with a z-score of 5.8, was the C-terminus of *S. cerevisiae* Swt1 (PDB: 4pqz), an RNA endonuclease that is a member of the Higher Eukaryotes and Prokaryotes Nucleotide binding protein (HEPN) superfamily, which includes nucleotide-hydrolyzing toxins [156,157]. The Swt1 HEPN domain matches four helices in pTox K28, from the alpha, gamma, and beta domains and postranslational cleavage of pTox would separate these helices, disrupting the predicted HEPN domain structure. Furthermore, the K28 alpha domain alone is responsible for toxicity and lacks the conserved consensus motifs required for hydrolysis of the phosphodiester backbone of nucleotides. Thus, it is unlikely that the structural match of K28 to HEPN domains is functionally relevant. Given the functional diversity of the remaining proteins identified by DALI and the low confidence of the K28 model, we concede that the structure and antifungal mechanism of K28 continue to remain enigmatic.

### 2.10 The K62 killer toxin family

The K62 killer toxin was first identified in *S. paradoxus* strain Q62.5 isolated from an English oak tree [101,113]. Based on its antifungal spectrum of activity against other killer yeasts, K62 was initially misclassified as a K1 toxin. Further investigations of K62 found that it was encoded by the satellite dsRNA named M62, and the genetic sequence of the novel K62 killer toxin was confirmed [100].

#### 2.10.1 The K62 killer toxin family: Domain organization and posttranslational modification

K62 is a 272-amino acid protein, shorter than most known *Saccharomyces* killer toxins, which average 320 amino acids in length. K62 shares no detectable amino acid sequence homology with any previously characterized killer toxins, and the secondary structure prediction exhibited a high proportion of beta strands (41.6%) and few alpha helices (4.1%). This secondary structure composition contrasted with other *Saccharomyces* killer toxins, which on average contain a higher proportion of alpha helices.

K62 has a signal peptidase cleavage site between amino acids 31 and 32, which is consistent with it being an extracellular killer toxin. However, unlike all other *Saccharomyces* killer toxins, the K62 sequence has only a single dibasic KR motif at residues 111 and 112. This motif is positioned on a solvent-exposed loop separating the N-terminal domain from the elongated C-terminal domain. This location suggests a potential Kex2 protease cleavage site, analogous to those found in other *Saccharomyces* killer toxins. However, unlike K1 and K28 killer toxins, such a cleavage would separate the N- and C-terminal domains due to the absence of cysteine residues in the N-terminal domain, which are required to form an interdomain disulfide bond. PSIPRED predicted a region between 4β and 3α that could interact with membranes and form pores, setting K62 apart from other killer toxins that predominantly have alpha helices that are predicted to be transmembrane and pore-forming.

#### 2.10.2 The K62 killer toxin family: Molecular modeling

The AlphaFold2 model of K62 predicts an elongated architecture (∼86 × 40 Å), characterized by extended beta sheets, which deviates substantially from the compact globular folds typical of other *Saccharomyces* killer toxins. The C-terminal domain consists of a five-stranded beta-sheet arrangement. In this configuration, beta strands 3β, 6-9β form a scaffold that is stabilized by three intramolecular disulfide bonds: one between C146-C271 linking the flexible loop between beta strands 3-4β to the C-terminal tail; a second between C201-C262 stabilizing beta strands 5β and 9β; and a third across a loop at the end of beta strand 6β (C227-C237). A second sheet is formed by strands 4β and 5β that flanks a flexible loop containing alpha helix 3α. This loop exhibits an alternating sequence of hydrophobic and polar residues enriched with serine and threonine, residues often implicated in promoting oligomer assembly and facilitating membrane engagement and pore formation in toxins.

#### 2.10.3 The K62 killer toxin family: Mechanistic insights

Analysis of the predicted tertiary structure of K62 using DALI yielded a top hit of parasporin-2 from the bacterium *Bacillus thuringiensis* with a z-score of 5.2 (PDB: 2ZTB). Parasporin-2 is an aerolysin family protein, a class of beta-barrel, pore-forming toxins that are bacterial virulence factors and highly toxic to eukaryotic cells [158–160]. Aerolysin toxins are thought to contribute to hemorrhaging and tissue necrosis in fish and small ruminants, with lethal dose 50% (LD-50) values as low as 100 ng/kg [159,161,162]. Moreover, parasporin-2 can recognize, permeabilize, and kill human cells [163]. The calculated RMSD of the aerolysin core domain of parasporin-2 compared to K62 was 6.9 Å, indicating structural homology despite only 16.8% and 32.1% amino acid identity and similarity. Several other K62 structural homologs from the aerolysin family of pore-forming proteins and toxins were also identified by DALI analysis. These homologs included members of a class of aerolysins found in bony fish and lamprey named natterins, which are theorized to play a role in innate immunity [164]. The natterin-like protein Dln1 from *Danio rerio* (zebrafish, z-score 4.3) binds mannans, including those of the fungal cell wall. Dln1 also undergoes pH-dependent oligomerization, with a low optimal pH similar to killer toxins [164]. Other K62 homologs were the HA3 toxin from the bacterium *Clostridium botulinum* (z-score 4.2) [165] and the innate immune stimulator BmALP1 from *Bombina maxima* (Yunnan firebelly toad, z-score 3.8) [166] (Figure 12). Notably, the receptor-binding domain of parasporin-2 and other K62 homologs had no structural similarity to the modelled K62 N-terminal domain. Although K62 is the first aerolysin family protein found in the *Saccharomyces* genus of yeasts, it is not the first fungal aerolysin toxin. The monomeric structure and biological activity of a hemolytic lectin from *Laetiporus sulphureus* was previously identified as a novel fungal aerolysin family protein [167]. However, the structure of this conserved aerolysin domain is not a close homolog of K62 (RMSD 9.8 Å). Bioinformatics studies have identified 41 fungal sequences from 19 different species with homology to aerolysin family proteins, but none of these aerolysin toxins were sequence or structural homologs of K62 by PSI-BLAST or DALI analysis [168] .

**Figure 12.**
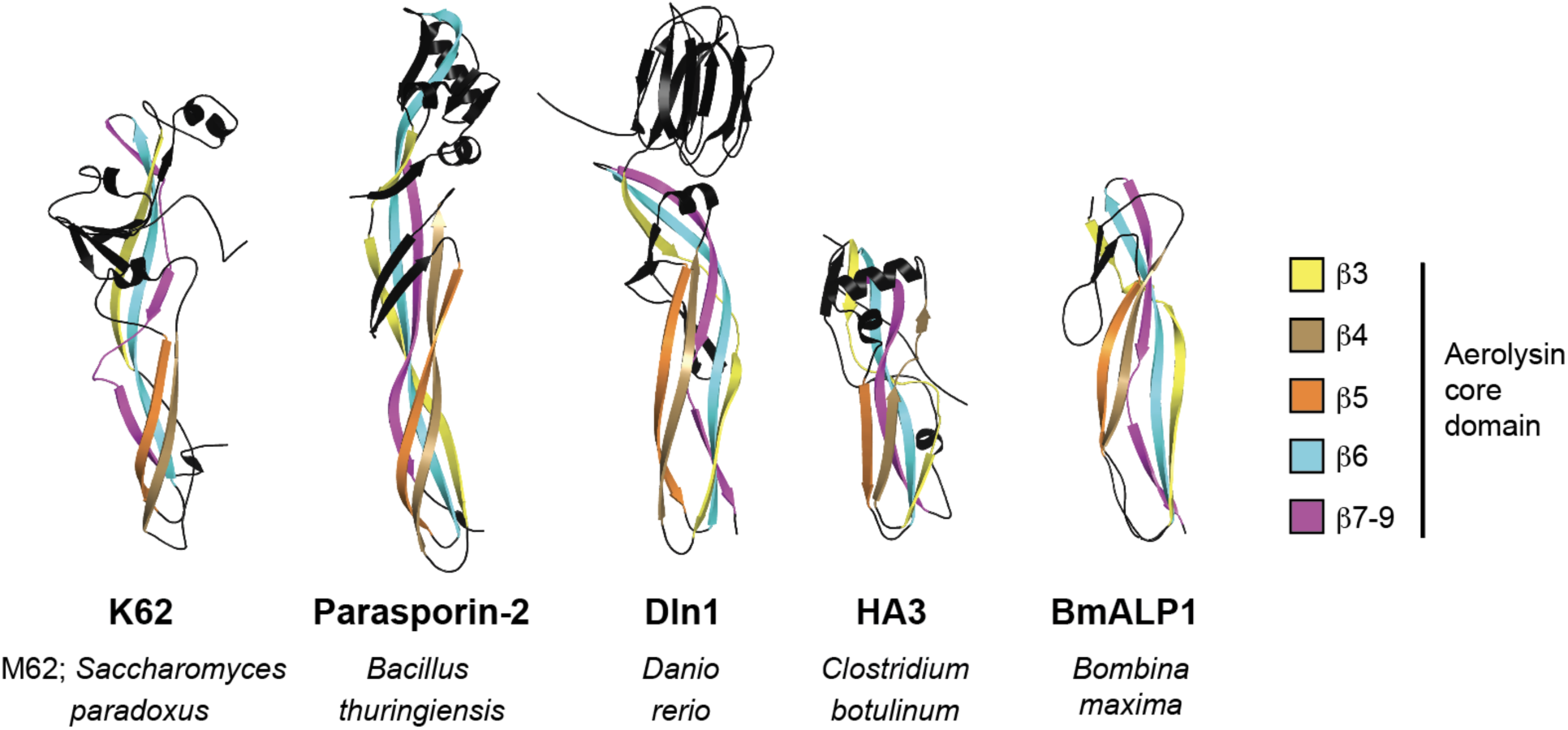
K62 structural homologs identified by DALI analysis. Tertiary structure models are colored by beta strands that contribute to the conserved five beta sheet motif of the aerolysin core domain in K62. The N-terminal receptor binding domain and insertion loop are colored black. PDB accession numbers for parasporin-2, Dln1, HA3, and BmALP1 are 2ZTB, 4ZNO, 2ZS6, and 6LH8, respectively.

Aerolysin toxins are identified by an elongated beta sheet structure and are typically composed of two domains. Firstly, an N-terminal receptor binding domain typically interacts with the target cell surface, providing specificity of the toxin to a cell type. The N-terminal domain tends to be more diverse within the aerolysin family of proteins, reflecting different cell-surface targets, such as carbohydrates and lipids. Secondly, the more conserved C-terminal aerolysin domain is responsible for beta-barrel pore formation that penetrates membranes, causing ion leakage and cell death.

Oligomerization of K62 monomers to form a pre-pore structure would be an essential first step before membrane attack by pore formation. Based on known aerolysin family pre-pore structures, K62 strands 4β and 5β would oligomerize with the same beta strands of neighboring monomers in an alternating pattern and form an inner ring, while beta strands 3β, 6-9β form an outer ring [169]. After pre-pore formation, beta strands 4β and 5β and the insertion loop containing 3α would penetrate the membrane, forming the extended beta barrel pore [164,169–172]. The enrichment and alternating pattern of polar serine (**S**) and threonine (**T**) residues with hydrophobic amino acids from position 160-196 in K62 (160-L**S**W**S**Y**T**Y**T**W**S**YDV**S**IGI**S**WEVI**S**A**S**VDY**S**I**S**Q**S**L**S**Y**S**-196) is characteristic of the membrane insertion loop of aerolysin-family toxins. The putative insertion loop of K62 also correlates with a predicted membrane-interacting region that was identified by PSIPRED (Figure 12A). Based on these similarities in sequence and structure, it is predicted that K62 is an ionophoric toxin that, after oligomerization, forms a beta-barrel pore to penetrate target membranes. More recent and thorough *in silico* and empirical investigation of K62 structure and function support this prediction.

Despite similarities to aerolysin toxins, K62 has several unique structural features. Specifically, the possible cleavage of the N-terminal domain from the aerolysin core domain by Kex proteases at a dibasic motif. This would represent a departure from the canonical aerolysin family toxins, where the N-terminal domain typically contributes to membrane and receptor recognition. Removal of the K62 N-terminal domain would be similar to aerolysins that have a minimal N-terminal domain, such as monalysin, which has an aerolysin core domain that is necessary and sufficient for membrane targeting and pore formation [173]. It is therefore plausible that the K62 core domain alone retains toxicity and is independent of the N-terminal region. The N-terminal domain may remain associated with the aerolysin core domain through non-covalent interactions, similar to the SMKT killer toxin. Another possibility is that immature K62 pTox may prevent the unwanted toxicity of the mature K62 before it is exported from the cell. The removal of the N-terminal domain by Kex cleavage could activate the toxin, analogous to the proteolytic processing required to activate aerolysin toxins [174,175]. Another unique feature of K62 that differs from the well-studied aerolysin proteins is the presence of disulfide bonds, which are common in killer toxins and in K62 are essential for toxicity.

Despite differences from known aerolysin toxins, confident structural predictions of K62 and its homology to parasporin-2 indicate it is a new member of the aerolysin family of toxins. The potency of aerolysin toxins and the presence of K62 and its homologs in fungal species relevant to plant, animal, and human health, as well as industrial fermentations, justify further study of their toxicity to cells of higher eukaryotes.

## 3. Conclusions

This study presents the most comprehensive structural and functional analysis of *Saccharomyces* killer toxins to date, integrating decades of empirical research with recent advances in tertiary structure prediction and MD simulations. These models will benefit the study of *Saccharomyces* killer toxins, advancing the functional understanding of these potent antifungal proteins and also increasing our knowledge of thousands of sequence and structural homologs. Using *S. cerevisiae* to study these toxins will help to achieve a greater understanding of how they interact with and intoxicate fungal cells and their possible future application against pathogenic and spoilage fungi. Furthermore, the homology of killer toxins to bacterial virulence factors and other cytotoxic proteins warrants investigation of their potential contribution to diseases of humans, plants, and animals.

Creating tertiary structure models of all *Saccharomyces* killer toxins revealed unexpected structural similarities between the seven members of the K1, K2, K74, and K45 killer toxin families that shared little to no amino acid sequence similarity (Figure S7). The shared core tertiary structure, domain organization, and patterns of postranslational modification allowed their designation as the K1 superfamily. All of these toxins included a central hydrophobic alpha helix buried within an α/β sandwich. This most likely indicates a common ionophoric mechanism of antifungal activity based on the proposed pore-forming antifungal mechanism of K1. Structural homology to SMKT also supports a mechanism of membrane attack whereby the toxic alpha stably associates with the membranes of intoxicated cells. However, direct evidence for the structure or assembly of killer toxin pores for any of these killer toxins has remained elusive.

A similar α/β sandwich organization was also determined for the Klus family, which had a closer structural similarity to SMKT than K1 superfamily toxins. Thus, it is possible to speculate that all of the aforementioned toxins have a similar mechanism of membrane attack by pore formation. The Klus family also has structural homology to toxic lectins form beta barrel pores, toxic filaments and the inhibition of calcium channels. These mechanisms differ from the proposed cation-specific alpha helical membrane pores formed by K1, but the exact role of this central helix has never been determined. Similarly, toxins from the Klus family exhibit structural similarity to well-studied lectin proteins from fungi and plants that bind specific carbohydrate moieties on cell surfaces. This homology is consistent with their proposed mode of action involving cell surface recognition and suggests that Klus family toxins may have evolved from carbohydrate-binding domains that were adapted for antifungal activity.

Equally surprising was the identification of K62 as a structural outlier among *Saccharomyces* killer toxins. Structural modeling placed K62 within the aerolysin-like pore-forming toxin family, a group more commonly associated with bacterial toxins. This discovery significantly expands the functional and structural diversity of killer toxins within the fungi and opens new questions regarding their mechanisms of action, evolutionary origins, and potential biotechnological applications.

This approach also highlighted limitations in structure prediction for certain toxins. Models of K28 and K74 were of low confidence and unstable during MD simulations. These challenges may reflect unique features not readily captured by current predictive algorithms. These cases emphasize the need for empirical validation and careful interpretation when using structural predictions algorithms, especially for proteins lacking close structural homologs or exhibiting unique processing and function.

Beyond the structural findings, our work also highlights the need for a more consistent and informative nomenclature for fungal killer toxins. Historically, toxins have been named somewhat arbitrarily, based on discovery order (e.g., K1, K2), source organism or strain (e.g., K21, K45, K74), or biological activity (e.g., KHR: killer of heat-resistant yeasts). Moreover, dsRNA-encoded killer toxins have not followed the standardized gene naming conventions used in *S. cerevisiae* and other model fungi. Given the growing number of homologous killer toxin sequences and their clear structural grouping into families, we propose a nomenclature informed by both sequence homology that can be further validated by structural modeling. For instance, we previously designated K1 homologs as “K1-like Killer Toxins” (KKT). We now propose gene names for the remaining families: KTT (K2; Killer Toxin 2), KTF (K45; Killer Toxin 45), KTS (K74; Killer Toxin 74), KTN (Klus; Killer Toxin NaCl-mediated/Klus family), and KTA (K62; Killer Toxin Aerolysin). These designations retain a connection to the first-discovered member of each family and reflect their predicted structural and mechanistic classifications.

In this manuscript, we refer to a representative K74 homolog as KTS1^Cmal^ (gene: *KTS1*), indicating that it belongs to the K74 family and was identified in *Cadophora malorum*. The superscript provides essential taxonomic context for comparing homologs across species. This standardized naming approach offers clarity, reduces redundancy, and allows researchers to draw meaningful functional comparisons across species. It also facilitates a more systematic exploration of killer toxin biology, enabling insights gained from well-characterized members, like K1 and K2, to accelerate our understanding of more recently discovered or poorly understood toxins.

In summary, this work establishes a framework for the structural classification and functional study of killer toxins in *Saccharomyces* and beyond. It reveals unexpected evolutionary and mechanistic relationships, proposes a unifying nomenclature, and sets the stage for future experimental studies into a diverse and potent group of fungal antifungal proteins.

## Methods

### Identification of Killer Toxin Homologs

To identify homologs of killer toxins, publicly available databases were searched using NCBI BLAST. Protein sequences of the *Saccharomyces* killer toxins, K1, K1L, K2, K21, K28, K45, K62, K74, KHR, KHS, and Klus were used as queries for position-specific iterated BLAST (PSI-BLAST) searches against the NCBI database. Each PSI-BLAST analysis was conducted until convergence was reached or 1,000 related sequences were retrieved. Results (4,437) were filtered to exclude proteins that were less than 75%, or more than 150% of the length of the query sequence, which removed 1,703 potential nonfunctional genes or proteins with significant departure from the length of known killer toxins. PSI-BLAST of K62, KHR, and Klus were the only and 663 homologs, respectively.

### AlphaFold2 and 3

All predicted structures presented in this study were generated using AlphaFold2 (version 2.2.0) with default parameters. Structure prediction was performed via singularity.py script on the University of Idaho’s Research Computing and Data Services platform. For monomer predictions, five models were generated for each sequence with amber relaxation and pLDDT was used to select the top ranked model for further analysis. The same protocol was used for each multimer prediction except that 25 models were generated instead of 5. All structures predicted by AlphaFold3 were done through the AlphaFold3 Server https://alphafoldserver.com. Sequences were acquired from NCBI Genbank, pasted into the ‘sequence’ field in AlphaFold Server, and submitted.

### Molecular Dynamics

AlphaFold2 results were prepared for molecular dynamics with a dodecahedral bounding box, solvated with SPC water, and solvent molecules were replaced with ions using Monte Carlo placement to neutralize the system. Each system was energy minimized using the steepest descent until the maximum force within the system reached 1000 kJ/mol/nm. NVT equilibration was performed on each energy-minimized system for 100 ps and temperature set to 300 K. Each system was then subjected to NPT equilibration for another 100 ps and pressure set to 1 atm. Production MD was then performed for 1 us. The GROMACS v2024.2 molecular dynamics engine, the SPC water model and the AMBER99-ILDN forcefield parameters were used for all simulations. The V-rescale thermostat was used for equilibration and production. The C-rescale barostat was used for NPT equilibration and production. All equilibration steps were performed with the heavy atoms of the protein restrained. For all simulations, the LINCS algorithm was used to enforce proper bond lengths with hydrogen. Electrostatics were calculated with Particle Mesh Ewald. Both electrostatic and Van der Waals cutoff distances were set to 1 nm. All simulations used a timestep of 2 fs. Analysis of trajectories was performed using GROMACS package tools. RMSD over time methods used backbone atoms. The clustering method used was pairwise Root Mean Squared Deviation (RMSD) and was performed until the lowest cutoff that resulted in 10 bins or less was selected.

### FoldX

FoldX version 5.0 was used to calculate ΔΔG of stability of point mutations of cysteine to alanine on the amber relaxed top ranked model. Prior to mutation and ΔΔG calculation the model was subjected to FoldX RepairPDB six times which has been shown to improve accuracy of ΔΔG calculation. PositionScan command was used to create mutant structures and calculate energy.

### Software

SignalP https://dtu.biolib.com/SignalP-6 PyMOL 2.6 https://www.pymol.org/

PSIPRED - Add a note on the use of MEMSTAT https://bioinf.cs.ucl.ac.uk/psipred

