## Supplementary figures and tables for "A Comprehensive Structural and Functional Analysis of *Saccharomyces* Killer Toxins"

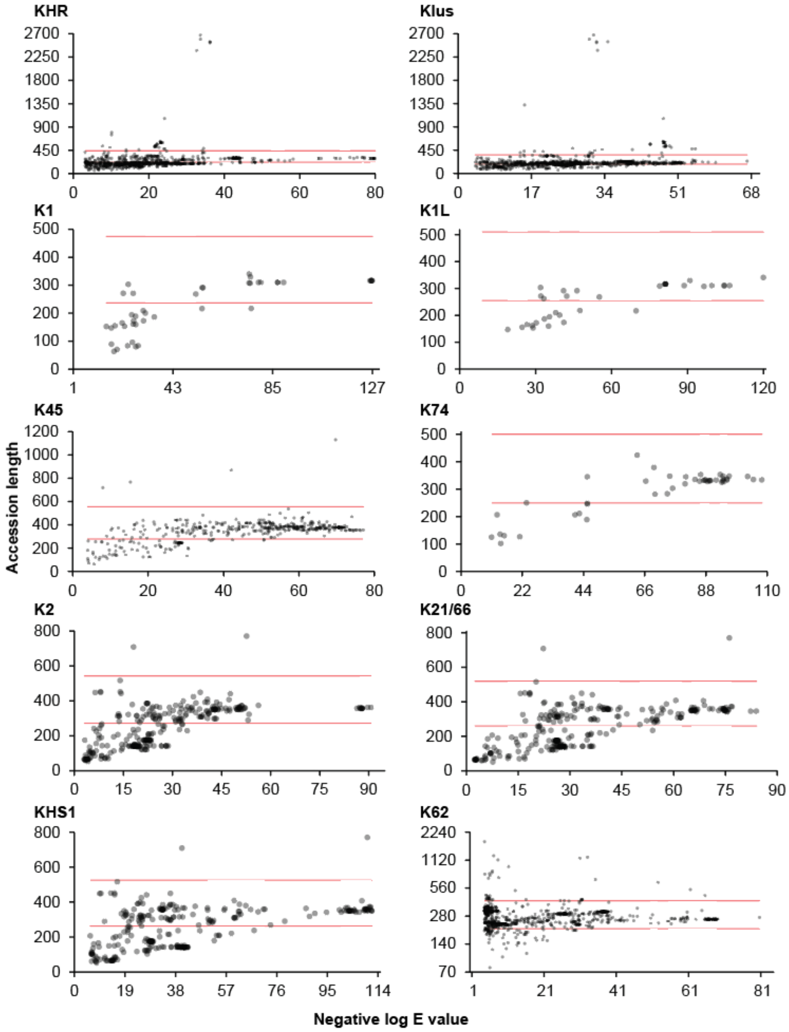


**Figure S1. Graphical PSI-BLAST results.** Red lines indicate size cutoffs.

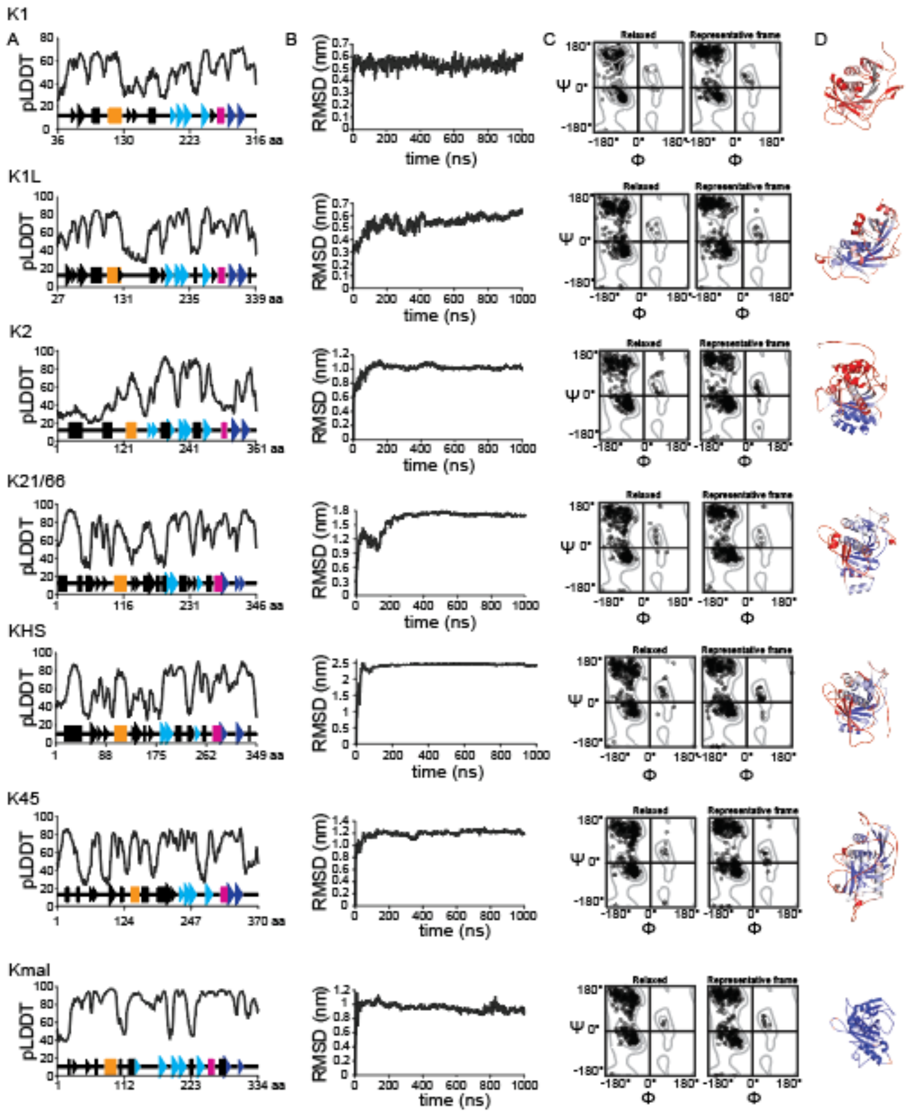

K1
A
B
C
D
K1L
K2
K21/66
KHS
K45
Kmal
pLDDT
RMSD (nm)
time (ns)
Relaxed
Representative frame
Ψ
Φ
aa

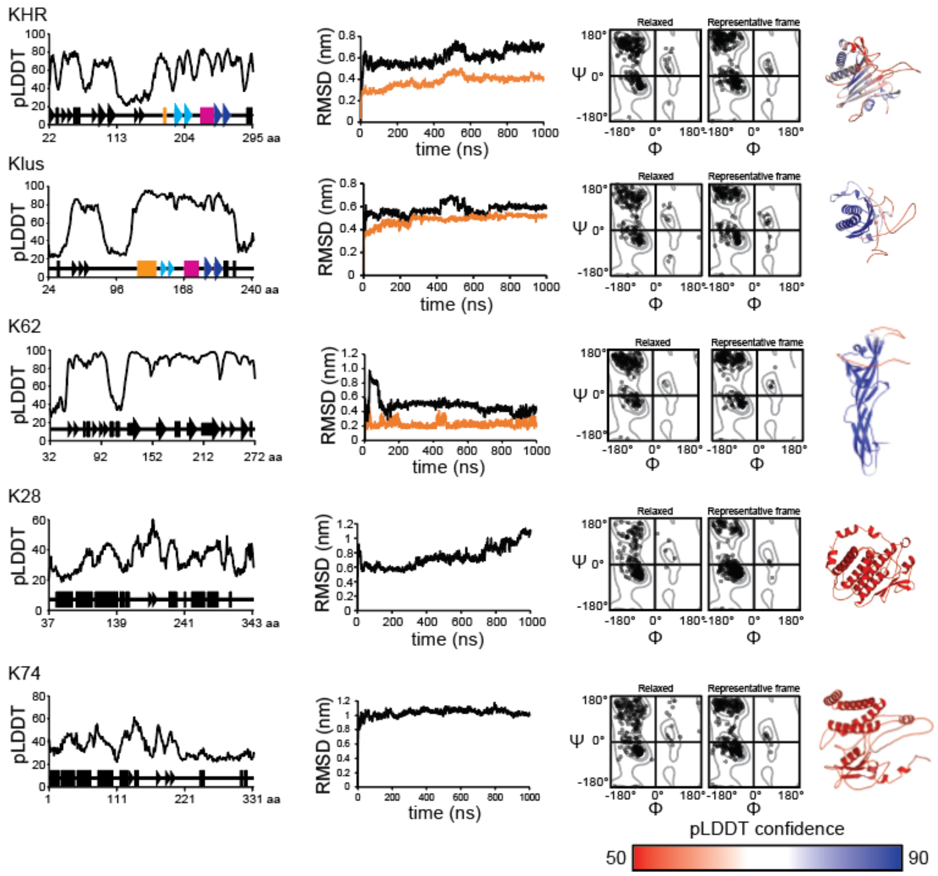


**Figure S2. AlphaFold confidence scores and RMSD trajectories of killer toxin tertiary structure models.** (A) Predicted local distance difference test (pLDDT) scores from AlphaFold2 models of all *Saccharomyces* killer toxins showing the confidence in the top-ranked model. (B) Full protein RMSD over 1 µs molecular dynamics simulation. GROMACS was used to generate alignments of each snapshot to the structure at 0 ns. Orange RMSD traces in KHR, Klus, and K62 have 111 to 161 residues removed, 15 N-terminal residues removed and 112 N-terminal residues removed, demonstrating the RMSD fluctuations are due to N-terminal flexibility. (C) Ramachandran plots of general residues (non-proline/glycine) generated by SWISS structure assessment tool before (AlphaFold2's amber relaxed output) and after MD simulation. (D) Top relaxed AlphaFold2 models colored by pLDDT confidence.

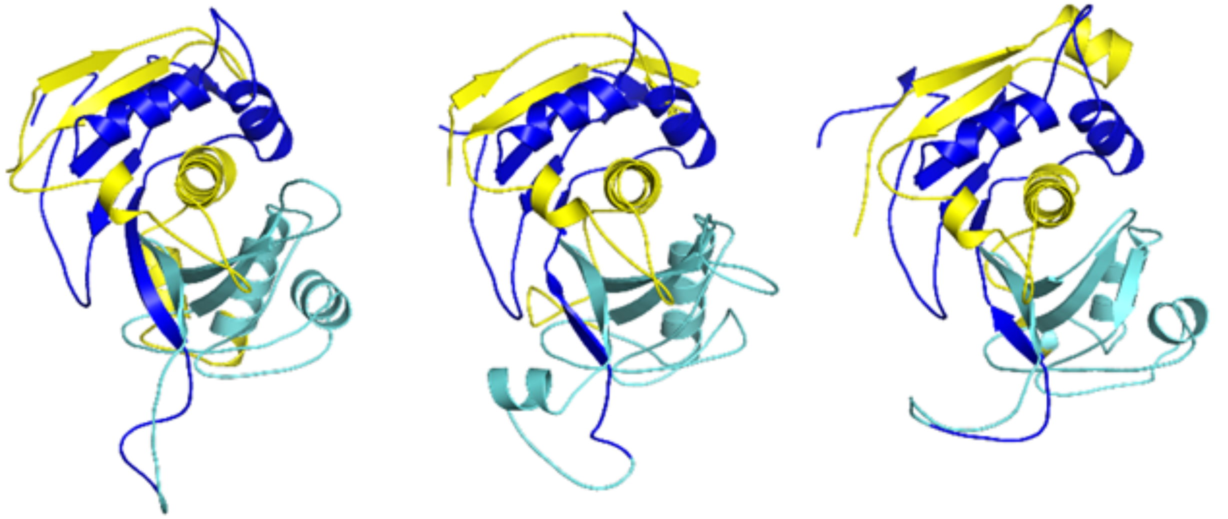

**Figure S3. AlphaFold3 predicted structures of representative K74 homologs.** Left to right: Accession: KAI6714648.1 from *Diplocarpon mali*, accession: CZR64371.1 from *Phialocephala subalpin*, and accession: OJJ75601.1 from *Aspergillus brasiliensis*. Structures colored by predicted alpha domain (yellow), gamma domain (cyan), and beta domain (blue) based on dibasic motif.

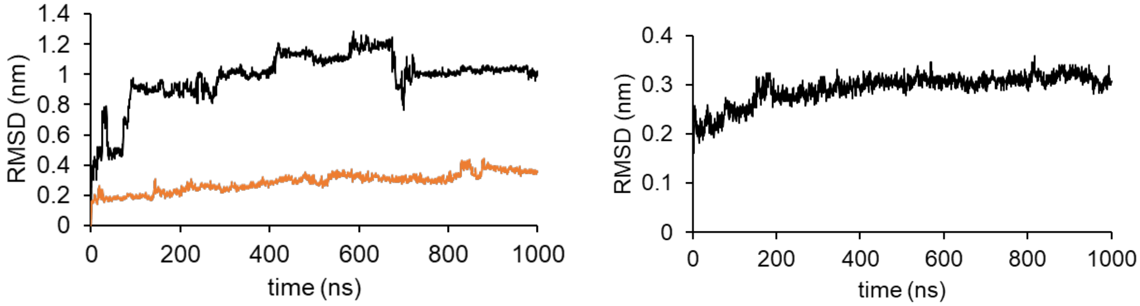


**Figure S4. Molecular dynamics simulations of mature Klus and KHS.** RMSD of backbone atoms of mature Klus (left) and mature KHR (right) compared to the starting conformation over 1 μs MD simulation. The orange line on Klus is the backbone atom RMSD minus the 30 C-terminal residues of the beta domain.

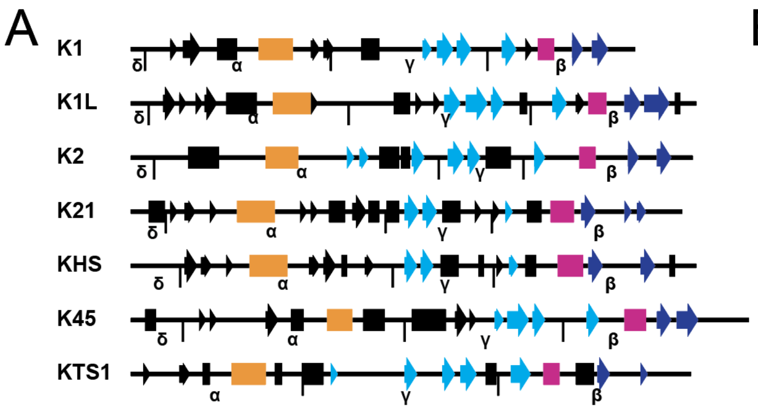


**Figure S6. Linear representation of K1 superfamily killer toxins and tertiary structure honologs colored by conserved secondary structure features.** Linear representation of proteins representing (A) killer toxins and (B) pilins, pseudopilins, and SMKT. Secondary structure elements that are conserved across structures are colored to highlight similarities including the central hydrophobic helix (orange) and a second conserved helix (magenta) that are positioned before and after beta strands (cyan) ending in C-terminal beta strands (dark blue). Structural features colored black are not considered to be conserved between different proteins. Alpha helices and beta sheets shown as rectangles and arrows, respectively. PDB accession numbers are included after gene names for the pilins, pseudopilins, and SMKT.

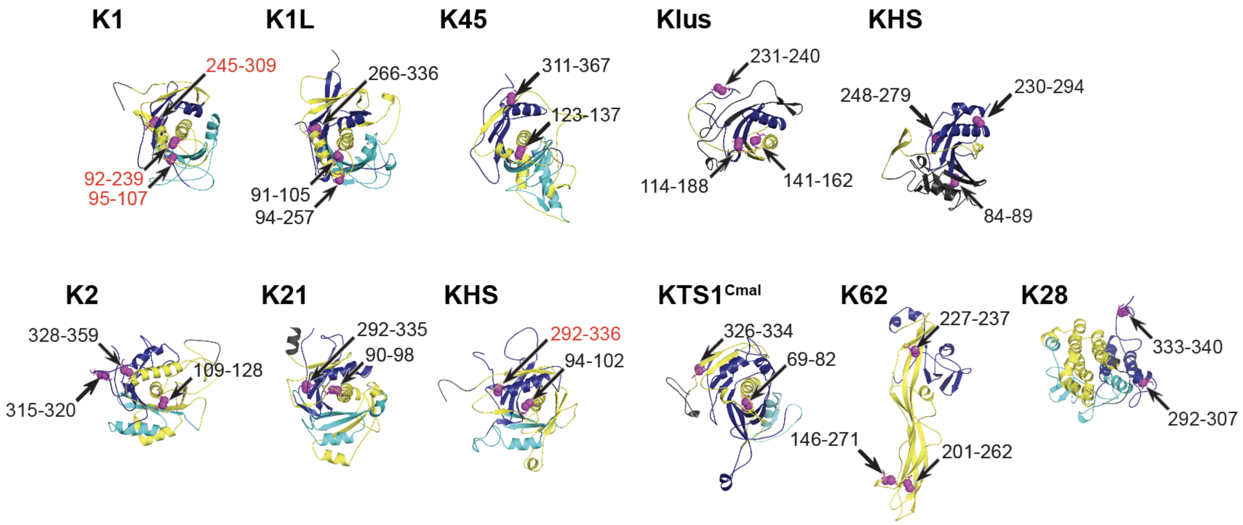


**Figure S7. Relative positioning of cysteine pairs in all killer toxin tertiary structure models.** Cartoon representations of pTox tertiary structures colored by their delta (black), alpha (yellow), gamma (cyan), and beta domains (dark blue). Cysteine pairs are represented by adjacent magenta spheres and are labelled by amino acid numbers (unpaired cysteine residues are omitted for clarity).

### Supplementary Tables

| Toxin | PDB ID | Residues |
|---|---|---|
| SMKT | 1KVD, 1KVE | 222 |
| KP4 | 1KPT | 127 |

| | | |
|---|---|---|
| WKT | 1WKT | 125 |
| KP6 | 1KP6, 4GVB | 219 |
| Zt-KP6-1 | 6QPK, 9GWD | 162 |
| Zt-KP4-1 | 8ACX | 96 |

*Table S1. Fungal killer toxins with experimentally determined tertiary structures*

| | *Killer toxin* | | | | | | | | | | |
|---|---|---|---|---|---|---|---|---|---|---|---|
| | *K1* | *K1L* | *K2* | *K21* | *KHS* | *K28* | *K45* | *K62* | *K74* | *Klus* | *KHR* |
| *Total* | 43 | 44 | 287 | 283 | 282 | 3 | 490 | 1000 | 45 | 1000 | 1000 |
| *Hits* | 20 | 21 | 145 | 145 | 145 | 3 | 354 | 883 | 34 | 643 | 322 |
| *Long* | 0 | 0 | 2 | 2 | 2 | 0 | 4 | 43 | 0 | 44 | 38 |
| *Short* | 23 | 23 | 140 | 136 | 135 | 0 | 92 | 74 | 11 | 313 | 640 |
| *Actinomycetes* | 0 | 0 | 1 | 1 | 1 | 0 | 0 | 0 | 0 | 0 | 0 |
| *Agaricomycetes* | 0 | 0 | 0 | 0 | 0 | 0 | 17 | 0 | 0 | 27 | 5 |
| *Alphaproteobacteria* | 0 | 0 | 0 | 0 | 0 | 0 | 0 | 0 | 0 | 0 | 1 |
| *Bacilli* | 0 | 0 | 2 | 2 | 2 | 0 | 0 | 36 | 0 | 0 | 0 |
| *Chytridiomycetes* | 0 | 0 | 0 | 0 | 0 | 0 | 0 | 3 | 0 | 0 | 0 |
| *Clostridia* | 0 | 0 | 0 | 0 | 0 | 0 | 0 | 1 | 0 | 0 | 0 |
| *Dipodascomycetes* | 0 | 0 | 0 | 0 | 0 | 0 | 3 | 0 | 2 | 0 | 0 |
| *Dothideomycetes* | 0 | 0 | 0 | 0 | 0 | 0 | 67 | 49 | 0 | 14 | 1 |
| *dsRNA* | 6 | 6 | 12 | 12 | 12 | 3 | 1 | 1 | 1 | 1 | 1 |
| *Eurotiomycetes* | 0 | 0 | 0 | 0 | 0 | 0 | 96 | 21 | 13 | 209 | 48 |
| *Exobasidiomycetes* | 0 | 0 | 0 | 0 | 0 | 0 | 0 | 1 | 0 | 1 | 0 |
| *Lecanoromycetes* | 0 | 0 | 0 | 0 | 0 | 0 | 0 | 0 | 0 | 10 | 1 |
| *Leotiomycetes* | 0 | 0 | 7 | 8 | 8 | 0 | 55 | 23 | 15 | 15 | 1 |
| *Lipomycetes* | 0 | 0 | 1 | 1 | 1 | 0 | 0 | 2 | 0 | 0 | 0 |
| *Lycopodiopsida* | 0 | 0 | 0 | 0 | 0 | 0 | 0 | 1 | 0 | 0 | 0 |
| *Magnoliopsida* | 0 | 0 | 0 | 0 | 0 | 0 | 1 | 0 | 0 | 0 | 0 |
| *Myxococcia* | 0 | 0 | 0 | 0 | 0 | 0 | 0 | 0 | 1 | 0 | 0 |
| *Pichiomycetes* | 3 | 3 | 19 | 19 | 19 | 0 | 6 | 65 | 0 | 45 | 29 |
| *Polypodiopsida* | 0 | 0 | 0 | 0 | 0 | 0 | 0 | 3 | 0 | 0 | 0 |
| *Saccharomycetes* | 12 | 12 | 102 | 101 | 99 | 0 | 18 | 116 | 2 | 88 | 199 |
| *Sordariomycetes* | 0 | 0 | 1 | 1 | 1 | 0 | 90 | 553 | 0 | 246 | 34 |
| *Tremellomycetes* | 0 | 0 | 0 | 0 | 0 | 0 | 0 | 8 | 0 | 1 | 0 |
| *Wallemiomycetes* | 0 | 0 | 0 | 0 | 0 | 0 | 0 | 0 | 0 | 6 | 2 |

*Table S2. Results of bioinformatics. Hits is all remaining accessions post size separation. legend needed.*

| | **pLDDT** | | | **Clustered** | | **Ramachandran favored** | | **Outliers** | |
|---|---|---|---|---|---|---|---|---|---|
| **Toxin** | **Global** | **Min** | **Max** | **%alpha** | **%beta** | **relaxed** | **rep frame** | **relaxed** | **rep frame** |
| K1 | 51.35 | 25.24 | 72.3 | 17.50% | 26.07% | 87.77% | 91.70% | 2.88% | 0.83% |

| | | | | | | | | | |
|---|---|---|---|---|---|---|---|---|---|
| K1L | 62.93 | 26.21 | 87.72 | 21.66% | 29.94% | 86.54% | 89.53% | 4.17% | 1.81% |
| K2 | 51.01 | 19.42 | 93.48 | 27.35% | 14.09% | 87.33% | 87.31% | 4.44% | 1.21% |
| K21/66 | 67.95 | 28.7 | 93.87 | 30.35% | 20.52% | 87.79% | 90.73% | 4.36% | 1.28% |
| KHS | 60.95 | 25.98 | 90.47 | 26.00% | 20.57% | 85.06% | 91.69% | 3.16% | 0.33% |
| K28 | 34.95 | 18.89 | 60.37 | 51.13% | 2.59% | 85.34% | 90.37% | 4.56% | 1.48% |
| K45 | 62.23 | 23.36 | 86.45 | 21.08% | 20.27% | 85.87% | 89.91% | 4.35% | 2.08% |
| K62 | 82.00 | 26.52 | 97.88 | 10.79% | 41.91% | 92.47% | 94.04% | 1.26% | 0.46% |
| K74 | 35.05 | 22.1 | 61.42 | 28.83% | 6.91% | 83.69% | 89.33% | 4.53% | 1.00% |
| Klus | 65.34 | 21.66 | 95.68 | 21.00% | 19.63% | 87.56% | 91.37% | 2.03% | 4.15% |
| KHR | 58.43 | 20.67 | 83.56 | 15.64% | 31.64% | 89.10% | 88.14% | 5.13% | 1.27% |
| KTS1 (AF3) | 80.60269461 | 39.12 | 97.65 | 20.1% | 17.1% | 94.58% | 90.82% | 0.60% | 1.31% |
| **Average** | **57.47** | **23.52** | **83.93** | **24.67%** | **21.28%** | **87.76%** | **90.41%** | **3.46%** | **1.43%** |

**Table S3. Confidence statistics of AlphaFold2 models.**

| Residue | Pos. | Mutation | delta delta G | Toxin activity on whole cells | Toxin Secretion | Cell wall receptor binding | Spheroplast killing | Immunity | Reference |
|---|---|---|---|---|---|---|---|---|---|
| G | 83 | D | 2.66 | <0.01 | 66 | + | - | - | [64] |
| C | 92 | Y | 9.13 | <0.02 | 65 | - | - | - | [64] |
| D | 101 | R | 0.51 | <0.05 | 3 | - | + | + | [64] |
| L | 114 | K | 2.39 | <0.01 | 59 | - | - | - | [64] |
| L | 115 | P | 6.44 | <0.01 | 41 | - | - | - | [64] |
| S | 124 | P | 0.01 | <0.01 | 21 | - | - | - | [64] |
| I | 129 | R | -0.15 | 10 | 80 | +/- | +/- | +/- | [64] |
| D | 140 | R | 0.37 | 10 | 80 | +/- | +/- | +/- | [64] |
| P | 237 | S | 3.57 | <0.1 | 68 | - | + | + | [64] |
| G | 264 | L | 3.86 | 75 | 75 | + | + | + | [64] |
| C | 312 | L | 6.32 | 50 | 57 | + | + | + | [64] |
| C | 92 | S | 3.19 | - | - | nd | - | + | [39] |

| | | | | | | | | | |
|---|---|---|---|---|---|---|---|---|---|
| C | 95 | S | 5.76 | - | - | nd | - | - | [39] |
| C | 107 | S | 4.23 | - | - | nd | - | - | [39] |
| C | 239 | S | 6.08 | - | nd | nd | - | + | [39] |
| C | 248 | S | 6.98 | - | + | nd | + | + | [39] |
| C | 312 | S | 6.27 | - | + | nd | - | + | [39] |
| T | 191 | P | 3.08 | 80 | 80* | nd | nd | + | [44] |
| R | 188 | A | 1.47 | 25 | nd | nd | nd | + | [44] |
| N | 181 | K | -0.35 | 40 | 40* | nd | nd | + | [44] |
| I | 151 | H | 1.64 | 100 | 100* | nd | nd | + | [44] |
| V | 85 | T | 0.62 | +/- | + | nd | nd | +/- | [62] |
| V | 116 | T | 2.15 | +/- | + | nd | nd | + | [62] |
| D | 101 | R | 0.517 | - | +/- | nd | + | + | [62] |

**Table S4. $\Delta\Delta G_{fold}$ values of mutations previously created in K1.**

| Toxin | SignalP | PSIPRED signal | Delta/ Alpha | Alpha/ Gamma | Gamma/ Beta | Other Kex cleavage sites |
|---|---|---|---|---|---|---|
| K1 | 36* | 33 | LLPR44* | VARR147* | VAKR234* | GERK100, YVKR188 |
| K1L | 26 | 12 | FDKR36 | PLKR147 | LTKRR248 | LNKK98, LTRR196 |
| K2 | none | none | VAVR63 | IVKR221 | VGKR268 | none |
| K21 | none | 40 | LHKR59 | LGKR181 | FTKR240 | none |
| KHS | none | 36 | LMRR63 | TIKR181 | FQKR237 | ATKR132 |
| Klus | 23 | 23 | | | | IIKR67, LAKR98, STVK167 |
| KHR | 21 | 16 | | | | ILKR77, VSKR130, FCKKK141, ATKR183 |
| K45 | none | 27 | IHRR56 | GFKKR179 | RAKR267 | FAKR97 |
| KTS1 | 23 | 38 | XXX | NNKR116 | LHRR156 | none |
| K74 | none | 38 | | | | LHKK102, WWKR110, TSRKR171, TVKR220 |
| K62 | 31 | 23 | n/a | n/a | n/a | LTKR112 |
| K28 | 36* | n/a | LEER49* | IQSR149* | LQKR245* | LYKR192 |
| SMKT | N/A | 35 | n/a | PVKR83* | PAAR145* | none |

*Table S5. Kex cleavage sites and domain boundary predictions for killer toxins. Cleavage sites are written as the four amino acids (in single letter nomenclature) occurring on the N-terminal side of the cleavage point. The numbering represents the number of the last amino acid in the sequence of four. n/a, not applicable. *indicates a site of postranslational cleavage that is supported by empirical evidence.*

| Toxin | Area (Å$^2$) | Solvent exposed (Å$^2$) | Buried (Å$^2$) | % buried | Helix sequence |
|---|---|---|---|---|---|
| K1 | 2026.2 | 328.2 | 1698.0 | 83.8 | CGKQTLALLVSIFVAVTSG |
| K1L | 2406.7 | 220.0 | 2186.8 | 90.9 | CFTAVSEVLKMSIMNAVKISI |
| K2 | 1730.0 | 77.8 | 1652.2 | 95.5 | PTLCAGAYVIGAMS |
| K21/66 | 2181.2 | 367.1 | 1814.1 | 83.2 | APLVNAILSTIVVSVAGGMAW |
| KHS | 2053.3 | 221.5 | 1831.8 | 89.2 | CAPIAGAVLATAAVIVAAVLV |
| Klus | 2336.5 | 950.5 | 1386.0 | 59.3 | YWDIADGVWQAGWDIYRATS |
| KHR | 501.0 | 120.1 | 380.5 | 76.0 | TDRI/ VYNLVSGITDRIKEAT* |
| K45 | 1301.0 | 142.9 | 1158.1 | 89.0 | DCVVGAVDTGVTLG |
| KTS1 | 1972.2 | 251.0 | 1721.2 | 87.3 | KAACQQSVLETGVCMGLSV |

*Table S6. Central helix buried surface area calculated using PyMol. KHR central helix calculations based on post MD clustered model.*

| Toxin | Validated disulfides | Disulfide bonds | Unpaired cysteines |
|---|---|---|---|
| K1 | C92-C239 | C92-C239, C95-C107, C245-C309 | |
| K1L | n/a | C91-C105, C94-C257, C266-C336, | C56, C94, C154, C159, C226 |
| K2 | n/a | C105-C128, C315-C320, C328-C359, | C25, C45, C300 |
| K21/66 | n/a | C90-C98, C292-C335 | C29 |
| KHS | n/a | C94-C102, C292-C335 | none |
| Klus | n/a | C114-C188, C141-C162, C231-C240 | none |
| Klus mature | n/a | C114-C188, C141-C162, C231-C240 | none |
| KHR | n/a | AF2 (C84-C89, C138-C151, C189-C219, C230-C294, C248-C279<br>MD (C84-C89, C230-C294, C248-C279) | AF2 (C143)<br>MD (C138, C143, C151, C189, C219) |
| KHR mature | n/a | C138-C294, C143-C230, C189-C219, C248-C279 | C151 |
| K45 | n/a | C123-C137, C311-C367 | C28 |
| KTS1 | n/a | C69-C82, C326-C334 | C92 |
| K74 | n/a | C70-C83, C284-C298 | C321, C329 |
| K62 | n/a | C146-C271, C201-C262, C227-C237 | none |
| K28 | C56-C333 | C292-C307, C333-C340 | C56 |

**Table S7. Predicted and experimentally determined disulfides and unpaired cysteines in killer toxins.**

| | | K1 | K1L | K2 | K21 | KHS | K45 | KTS1 | KHR | Klus | 2ret | 4noa | 3jyz | 7o6y | 5bw0 | 4d40 |
|---|---|---|---|---|---|---|---|---|---|---|---|---|---|---|---|---|
| AF2+ MD | K1 | 0.0 | | | | | | | | | | | | | | |
| | K1L | 5.0 | 0.0 | | | | | | | | | | | | | |
| | K2 | 11.1 | 6.1 | 0.0 | | | | | | | | | | | | |
| | K21 | 6.5 | 6.0 | 6.6 | 0.0 | | | | | | | | | | | |
| | KHS | 6.6 | 7.3 | 6.9 | 6.1 | 0.0 | | | | | | | | | | |
| | K45 | 5.7 | 5.8 | 5.1 | 5.1 | 6.4 | 0.0 | | | | | | | | | |
| | KTS1 | 5.7 | 6.5 | 8.1 | 6.3 | 6.2 | 6.8 | 0.0 | | | | | | | | |
| | KHR | 5.4 | 5.2 | 9.3 | 5.3 | 6.3 | 8.2 | 5.1 | 0.0 | | | | | | | |
| | Klus | 4.8 | 5.2 | 6.3 | 4.6 | 4.5 | 9.8 | 5.8 | 6.8 | 0.0 | | | | | | |
| PDB | 2ret | 6.3 | 7.5 | 5.7 | 5.3 | 5.3 | 7.5 | 5.5 | 5.5 | 9.3 | 0.0 | | | | | |
| | 4noa | 4.8 | 5.6 | 6.3 | 5.5 | 5.4 | 5.4 | 5.7 | 10.0 | 9.6 | 3.9 | 0.0 | | | | |
| | 3jyz | 4.8 | 5.8 | 6.0 | 5.4 | 5.4 | 6.2 | 5.5 | 5.6 | 7.8 | 6.1 | 4.3 | 0.0 | | | |
| | 7o5y | 9.2 | 6.9 | 7.5 | 5.9 | 6.5 | 6.7 | 6.2 | 5.4 | 4.5 | 4.7 | 4.3 | 6.3 | 0.0 | | |
| | 5bw0 | 6.5 | 9.2 | 5.9 | 5.6 | 5.1 | 7.2 | 5.8 | 8.4 | 8.8 | 2.3 | 4.8 | 6.5 | 4.0 | 0.0 | |
| | 4d40 | 6.4 | 8.7 | 8.2 | 6.4 | 5.1 | 6.2 | 6.0 | 5.7 | 9.3 | 4.9 | 2.5 | 3.9 | 4.9 | 4.9 | 0.0 |
| | 1kvd | 4.6 | 3.9 | 6.1 | 4.4 | 4.6 | 4.2 | 5.4 | 4.2 | 3.4 | 4.7 | 5.8 | 5.3 | 5.9 | 4.5 | 4.6 |

**Table S8. RMSD of ionophore killer toxins models to pilin crystal structures.** Measured using cealign command in pymol.

| | | Klus pptox | KHR pptox | Matue Klus | Mature KHR | SMKT | Kp4 | Gnk-2 | LDL | Zt- Kp4 | Y3 | VVA2 |
|---|---|---|---|---|---|---|---|---|---|---|---|---|
| AF1 + MD | Klus ppTox | 0.0 | | | | | | | | | | |
| | KHR ppTox | 6.8 | 0.0 | | | | | | | | | |
| | Mature Klus | 3.5 | 4.7 | 0.0 | | | | | | | | |
| | Mature KHR | 5.2 | 4.6 | 4.6 | 0.0 | | | | | | | |
| 1KVE | SMKT | 4.1 | 4.5 | 3.2 | 4.1 | 0.0 | | | | | | |
| 1KPT | Kp4 | 5.4 | 5.8 | 4.8 | 5.9 | 5.4 | 0.0 | | | | | |
| 3a2e | Gnk-2 | 4.5 | 4.5 | 4.7 | 4.9 | 4.5 | 5.4 | 0.0 | | | | |
| 4nds | LDL | 4.8 | 3.8 | 3.6 | 3.7 | 4.3 | 4.6 | 4.1 | 0.0 | | | |
| 8acx | Zt- kp4 | 4.9 | 5.0 | 4.7 | 6.2 | 8.2 | 5.0 | 5.5 | 3.8 | 0.0 | | |
| 5v6i | Y3 | 5.1 | 5.9 | 3.9 | 5.2 | 4.8 | 4.2 | 4.2 | 3.8 | 4.0 | 0.0 | |
| 1pp0 | VVA2 | 7.1 | 6.5 | 6.8 | 4.4 | 7.0 | 5.4 | 5.8 | 5.0 | 5.8 | 5.9 | |
| AF3 | SMKT ppTox | 3.6 | 4.2 | 4.8 | 6.5 | 4.2 | 5.7 | 6.1 | 7.9 | 4.7 | 7.7 | 6.4 |

Table S9. RMSD of Klus family killer toxins models to crystal structure matches from DALI. Measured using cealign command in pymol. AF1, AlphaFold2; AF3, AlphaFold3.
